# Spatially regulated protease activity in lymph nodes renders B cell follicles a sanctuary for retention of intact antigens

**DOI:** 10.1101/2021.11.15.468669

**Authors:** Aereas Aung, Ang Cui, Ava P. Soleimany, Maurice Bukenya, Heya Lee, Christopher A. Cottrell, Murillo Silva, Jesse D. Kirkpatrick, Justin R. Gregory, Parastoo Amlashi, Tanaka Remba, Leah M. Froehle, Shuhao Xiao, Wuhbet Abraham, Josetta Adams, Heikyung Suh, Phillip Huyett, Douglas S. Kwon, Nir Hacohen, William R. Schief, Sangeeta N. Bhatia, Darrell J. Irvine

## Abstract

The structural integrity of vaccine antigens is critical, as antigen degradation *in vivo* could eliminate neutralizing epitopes and create competing B cell responses against irrelevant breakdown products. Using FRET imaging and imaging zymography, we found that protease activity and antigen breakdown are spatially heterogeneous in lymph nodes. Following protein immunization, antigens are rapidly degraded in the subcapsular sinus, paracortex, and interfollicular regions of the tissue. By contrast, the follicles and follicular dendritic cell (FDC) networks exhibit low protease activity and antigen degradation rates. Immunization regimens targeting antigen rapidly to FDCs led to germinal centers (GCs) where responses to intact antigen were highly dominant, while traditional bolus immunizations led to weaker GC responses where more GC B cells bound to breakdown products than intact antigen. Thus, spatially-compartmentalized antigen proteolysis impacts humoral immunity and can be exploited to enhance vaccine-induced production of antibody responses against key pathogen structural epitopes.

## INTRODUCTION

Adaptive immune responses following infection or immunization are initiated when antigens are transported to lymph nodes by lymph or migrating immune cells. Humoral immune responses begin by B cells binding with their B cell receptors to cognate antigen, followed by the formation of germinal centers where B cells undergo proliferation and affinity maturation, leading to the production of high affinity antibodies against the target antigen (Pollard and Bijker, 2021; Shlomchik et al., 2019; Victora and Nussenzweig, 2012). It remains poorly understood what factors determine the makeup of the eventual affinity-matured polyclonal antibody response, though the precursor frequency of B cells specific for target epitopes, affinity of precursors for the antigen, antigen complexity, follicular helper T cell (T_FH_)-derived signals, and antibody feedback all contribute (Abbott et al., 2018; Bannard and Cyster, 2017; Dosenovic et al., 2018; Kuraoka et al., 2016; McNamara et al., 2020). In addition, the duration of antigen exposure and amount of antigen available to B cells plays an important role (Cirelli and Crotty, 2017; Tam et al., 2016).

An additional factor influencing the specificity of the antibody response following vaccination is the physical state of the antigen. To elicit protective responses, antigens need to preserve neutralizing epitopes that are faithful structural mimics of the target pathogen, which are often complex three-dimensional surfaces (Graham et al., 2019). Disruption of these epitopes has the potential not only to lower the availability of antigen for B cells with the capacity to produce neutralizing antibodies, but may also create distracting *de novo* epitopes that are irrelevant for protective immunity. It has been reported that model protein antigens can be rapidly proteolyzed as they reach the subcapsular sinus of lymph nodes; this antigen cleavage was linked to protease activity in serum/interstitial fluid (Catron et al., 2010). Intensive protein engineering efforts applied to viral envelope antigens have led to the generation of immunogens that may be stable in serum, but this may or may not render them resistant to all potential tissue-specific proteases. Such pathways of antigen breakdown might at least partially explain the substantial proportion of B cells that enter germinal center reactions but do not detectably bind to the immunizing antigen (Cirelli and Crotty, 2017; Kuraoka et al., 2016).

In contrast to potentially rapid antigen proteolysis in the SCS, several lines of evidence suggest that antigen trapped on dendrites of follicular dendritic cells (FDCs) may remain intact over extended time periods. For example, early studies showed that FDC-bound antigen recovered from lymph nodes after 12 weeks could be recognized by epitope-sensitive monoclonal antibodies and eluted in size exclusion chromatography in a manner suggesting gross antigen integrity (Tew et al., 1979). HIV virions deposited on FDCs in mice can be extracted from lymph nodes and functional viral particles recovered over several months, though the quantitative proportion of particles that are infective is not clear (Smith et al., 2001). FDCs have been shown to cyclically internalize and recycle trapped antigen, which may protect it from scavenging by B cells or extracellular degradation (Heesters et al., 2013), but whether this is the only or most important mechanism maintaining FDC-trapped antigen integrity remains unclear. These data collectively suggest that the follicles and the FDC networks in particular may be sites within lymph nodes where antigens are protected from degradation, while regions such as the sinuses may be areas of high proteolytic activity. However, to our knowledge the nature of protease activity in lymphoid organs has not been studied, and how antigen proteolysis impacts the immune response to vaccines is poorly understood.

We recently discovered that heavily glycosylated HIV Env or influenza immunogens prepared as multivalent protein nanoparticles are recognized by the serum protein mannose binding lectin (MBL) (Tokatlian et al., 2019). MBL binding to sugars present on these antigens subsequently triggers complement deposition on the nanoparticles. Following parenteral immunization, such particle immunogens initially arrive at the subcapsular sinus of lymph nodes (LNs), but within ∼48 hr, are transported to follicular dendritic cells (FDCs) in an MBL-and complement-dependent manner (Tokatlian et al., 2019). In these studies, we noticed that antigen outside of the FDC networks was rapidly cleared from LNs within a few days, while FDC-localized antigen persisted for weeks. While extrafollicular antigen loss could reflect clearance by lymph or internalization and intracellular degradation by lymph node cells, we hypothesized that extracellular protease activity could also play a role, though it remained unclear why antigen localized to FDC networks would be protected.

To shed light on these issues, we developed a FRET-based approach to track the integrity of antigens following subunit vaccine immunization, and analyzed the spatial pattern of protease expression and activity in lymph nodes. Unexpectedly, we found a pronounced spatial variation in protease activity, with high levels of antigen breakdown and protease expression in extrafollicular regions of mouse and human lymphoid tissues, but low levels of protease activity and high retention of antigen integrity over time within the FDC network of B cell follicles. Prompted by these findings, we evaluated the impact of antigen localization on the specificity of GC B cell responses, and found evidence that FDC-targeted protein immunizations achieve substantially greater proportions of antigen-specific B cell responses targeting conformationally intact epitopes compared to traditional bolus vaccination.

## RESULTS

### Monitoring antigen integrity through FRET analysis

To investigate the stability of vaccine antigens within lymph nodes (LNs) after immunization, we developed an approach labeling immunogens with paired small-molecule dyes capable of undergoing fluorescence resonance energy transfer (FRET), as a means to detect gross disruptions of antigen structure *in situ* in tissues. We hypothesized that antigen proteolysis would lead to separation of FRET donor and acceptor dyes, altering the magnitude of FRET signals induced in proportion to the degree of antigen degradation (**Figure 1A**). We employed acceptor photobleaching (Roszik et al., 2008), where the emission of the donor dye (Cy3) is measured before and after bleaching of the acceptor dye (Cy5) to monitor antigen integrity. This approach avoids complications of donor and acceptor excitation/emission crosstalk since only the donor emission is examined, and is independent of donor/acceptor dye concentrations and ratios on the protein of interest. As a test case for this approach, we labeled eOD-GT8 60mer (eOD-60mer), a ∼30 nm diameter protein nanoparticle (NP) presenting 60 copies of an HIV Env gp120 engineered outer domain (**Figure 1A**). eOD-60mer is a clinical vaccine candidate used as a priming immunogen to initiate CD4 binding site-directed broadly neutralizing antibodies (bnAbs) against HIV (Dosenovic et al., 2015; Jardine et al., 2013; Jardine et al., 2016; Jardine et al., 2015; Sok et al., 2016). This NP antigen was conjugated with amine-reactive Cy3 and Cy5 dyes at densities enabling substantial FRET on excitation of the donor Cy3 dye. The eOD-60mer NP accommodated labeling with at least 40 dyes per particle (∼20 Cy3 and ∼20 Cy5 dyes, eOD-60mer_40_) without disrupting binding of the bnAb VRC01 to the CD4 binding site epitope presented by the eOD-GT8 immunogen (**Figure S1A**). We previously showed that following a primary immunization, mannose binding lectin (MBL) present in serum binds to glycans densely displayed on the surface of the eOD-60mer, leading to subsequent complement deposition on the particles and trafficking of the antigen to FDCs within a few days (Tokatlian et al., 2019). Complement binding to labeled eOD-60mer incubated *in vitro* in mouse serum was also not statistically different from the unmodified immunogen for NPs labeled with up to at least 40 dyes per particle (**Figure S1B**). This degree of labeling represents modification of only 3.5% of the total amino acids of the particle immunogen. eOD-60mer_40_ coated onto glass coverslips showed a high level of FRET on excitation at 555 nm, but upon photobleaching of the Cy5 dye, emission in the Cy3 donor channel increased by 2-fold (**Figure S1C, G**). Such a change in Cy3 fluorescence was not observed for NPs labeled with only Cy3 or only Cy5, or when Cy3-labeled NPs were mixed with Cy5-labeled NPs and coated on coverslips together (**Figure S1D-G**). These changes in acceptor emission pre-and post-photobleaching can be quantified to determine the mean FRET efficiency *E* (see Methods). Based on these findings, we focused on NPs labeled with ∼40 dyes total for further studies.

**Figure 1.**
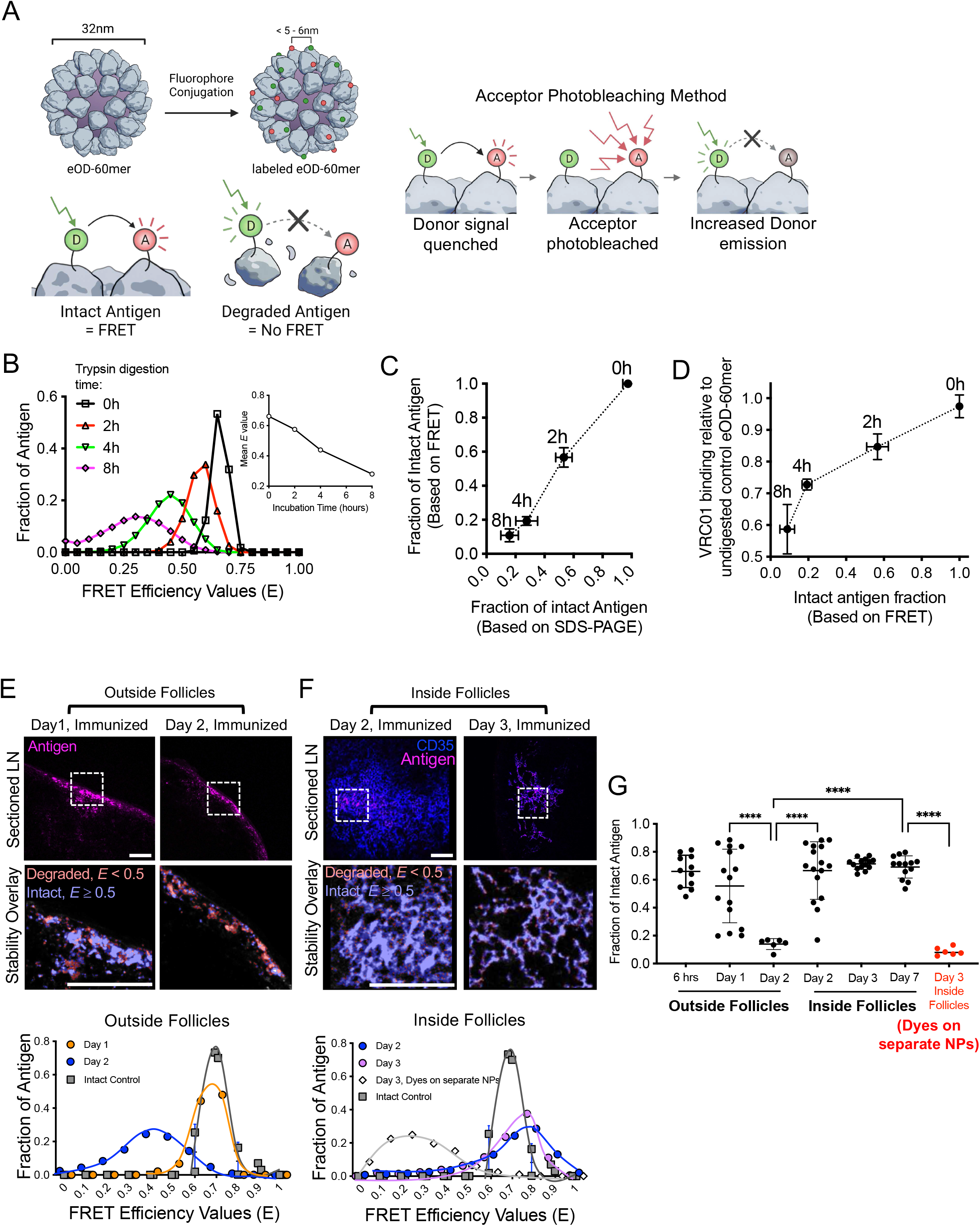
FRET-based analysis of immunogen integrity reveals spatially-variant antigen degradation in lymph nodes. (A) Schematic of FRET labeling of eOD-60mer protein nanoparticle immunogen and acceptor photobleaching strategy. (B-D) eOD-60mer_40_ particles were incubated with trypsin for indicated times and then coated on glass coverslips for FRET imaging. (B) Representative histograms of FRET Efficiency values, *E*, calculated from acceptor photobleaching. (C) Comparison of measured proportion of intact eOD-60mer_40_ as measured by FRET vs. SDS-PAGE. Shown are mean±s.d. (n = 3 samples/time point). (D) Comparison of VRC01 mAb binding to eOD6-mer_40_ vs. the FRET-measured proportion of intact antigen following trypsin digestion of the particles for varying times. Shown are mean±s.d. (n = 3 samples/time point). (E-G) C57BL/6J mice (n=4 animals/group) were immunized with 10 µg eOD-60mer_40_ and 5 µg saponin adjuvant. At indicated time points, LNs were harvested, flash frozen, and sectioned for confocal and FRET imaging. (E-F) Representative confocal images of eOD-60mer_40_ signal collected from a region near the subcapsular sinus (“Outside Follicles”, E) vs. within B cell follicles (“Inside Follicles”, F), showing antigen and anti-CD35 stain overlay (upper images) and higher magnification views (lower images, corresponding to dashed boxes in upper images) showing proportion of intact antigen determined by FRET imaging, illustrated using a false color scale binary image. Bottom graphs show the corresponding FRET Efficiency values from magnified regions comparing to fully intact control eOD-60mer_40_ imaged on coverslips prior to immunization. “D3, Dyes on separate NPs” in (F) indicate mice immunized with saponin adjuvant and 5 µg of two different eOD-60mers each labeled with only Cy3 or Cy5 dyes. All scale bars indicate 100 μm in length. (G) Mean fraction of intact eOD-60mer_40_ in LN at the indicated locations and times as determined by FRET imaging. Each point represents one region from one tissue section. Data collected from at least 8 tissue sections from 8 LNs. Shown are mean±s.d. (one-way ANOVA with post-hoc Tukey test for multiple comparisons, ****p ≤ 0.0001).

To determine how FRET efficiency relates to the structural integrity of labeled eOD-60mer particles, we calculated *E* distributions of intact or trypsin-digested eOD-60mer_40_. With increased digestion time, proteolyzed eOD-60mer_40_ showed the development of breakdown products by gel electrophoresis (**Figure S1H)**, which were accompanied by a shift in the distribution of measured *E* values for the particles, with mean *E* decreasing with time (**Figure 1B**). Importantly, the fraction of fully intact antigen as determined by FRET (calculated as the proportion of material with an *E* value overlapping the undigested control particle *E* distribution) was highly correlated to the fraction of intact antigen as measured by SDS-PAGE (**Figure 1C**). VRC01 binding to eOD-60mer_40_ captured on ELISA plates also decreased with increasing trypsin digestion time, demonstrating that the loss of FRET signal was not due simply to particle disassembly (**Figure 1D)**. Altogether, these data suggest that FRET signals from labeled 60mer NPs effectively report on the retention of intact immunogen structure.

### Antigens are rapidly proteolyzed at the subcapsular sinus but protected in follicles

We next used this FRET-based approach to analyze antigen integrity *in vivo* in mice following immunization. We previously reported that following primary immunization, eOD-60mer immunogens initially accumulate in the subcapsular sinus (SCS) of draining LNs, but over the course of several days are trafficked to the FDC network of follicles in an MBL- and complement-dependent manner, where they are highly enriched and retained for more than a week (Tokatlian et al., 2019). To assess the structural integrity of the NPs during this process, we immunized mice with FRET dye-labeled eOD-60mer_40_ and a saponin-based adjuvant, and at various time points after immunization, draining lymph nodes were extracted, flash frozen, and cryosectioned for acceptor photobleaching FRET signal analysis (**Figure S2A**). FRET efficiencies were converted to a measure of % intact antigen by comparing *E* values measured in the tissue to that of control fresh antigen coated on glass coverslips. As we observed previously, eOD-60mer could be detected in the SCS within a few hours post immunization, but over ∼48 hr, antigen levels in the SCS decreased, while steadily increasing amounts were detected on FDCs (**Figure S2B-C**). Unexpectedly, we observed location-dependent patterns of antigen degradation. eOD-60mer particles localized in the sinus or interfollicular regions of the LN exhibited a substantial loss in FRET efficiency between 24 and 48 hr (“Outside follicles”, **Figure 1E**). By contrast, eOD-60mer_40_ localized within the FDC network of follicles at the same timepoints exhibited high *E* values (**Figure 1F-G**, “Inside Follicles”). Quantification of the proportion of intact eOD-GT8 NPs from multiple follicles and multiple LNs revealed that a majority of eOD-60mer located outside of follicles was degraded within 48 hr, while over 70% of the antigen localized to the FDC network remained intact through at least 7 days (**Figure 1G**). Imaging of LNs from mice immunized with control mixtures of Cy3-eOD-60mer and Cy5-eOD-60mer showed negligible FRET signals for FDC-localized antigens, suggesting that sustained FRET signals observed for FDC-localized antigen are not due to inter-particle FRET (**Figure 1G**, “dyes on separate NPs”). As an orthogonal measure of antigen integrity, we also stained LN sections with VRC01 antibody to detect 60mer with an intact CD4 binding site epitope, and this approach revealed similar results: eOD-60mer in the SCS showed a high level of VRC01 staining at 6 and 24 hrs post immunization, but this staining was almost completely lost by 48 hrs for material outside of the FDC network (**Figure S2D-E**). By contrast, NP antigen localized to FDCs retained VRC01 staining even at 3 days post immunization (**Figure S2D-E**). Loss of VRC01 staining in extrafollicular sites was not due to an early antibody response blocking the epitope, because the same loss in VRC01 binding was observed in LNs from B cell receptor-transgenic MD4 mice where all B cells express an antigen receptor specific for an irrelevant antigen (Mason et al., 1992) (**Figure S2F**). Thus, both FRET and antibody staining analyses suggest that eOD-60mer particles are rapidly degraded as they enter the SCS and interfollicular regions, but antigen localized to dendrites of FDCs in follicles is protected for at least 7 days.

### Metalloproteinases are expressed in sinuses and stromal cells and contribute to rapid extrafollicular antigen degradation

We next sought to understand contributors to rapid extrafollicular antigen breakdown. A first possibility was that antigen was rapidly internalized and degraded by intracellular proteases outside the FDC network, whereas FDCs are known to rapidly recycle internalized antigens without degradation (Heesters et al., 2013). To assess this possibility, we immunized mice with dye-labeled eOD-60mer, and then stained non-permeabilized LN tissue sections with anti-dye and VRC01 antibodies to assess the proportion of intact extracellular antigen over time. This analysis revealed that antigen in extrafollicular regions was internalized over time, but 30% on average remained extracellular through 2 days post-immunization (**Figure 2A-B**), and extracellular antigen lost recognition by VRC01 over time in a manner very similar to the total antigen pool imaged by FRET (**Figure 2C**). We also examined the possibility that proteases in plasma may contribute to antigen degradation, but eOD-60mer incubated with plasma *in vitro* showed no loss in FRET signal (**Figure 2D**). Hence, a proportion of antigen outside B cell follicles remains extracellular in the lymph node for at least a few days, but a majority of this antigen is no longer fully intact by 48 hr.

**Figure 2.**
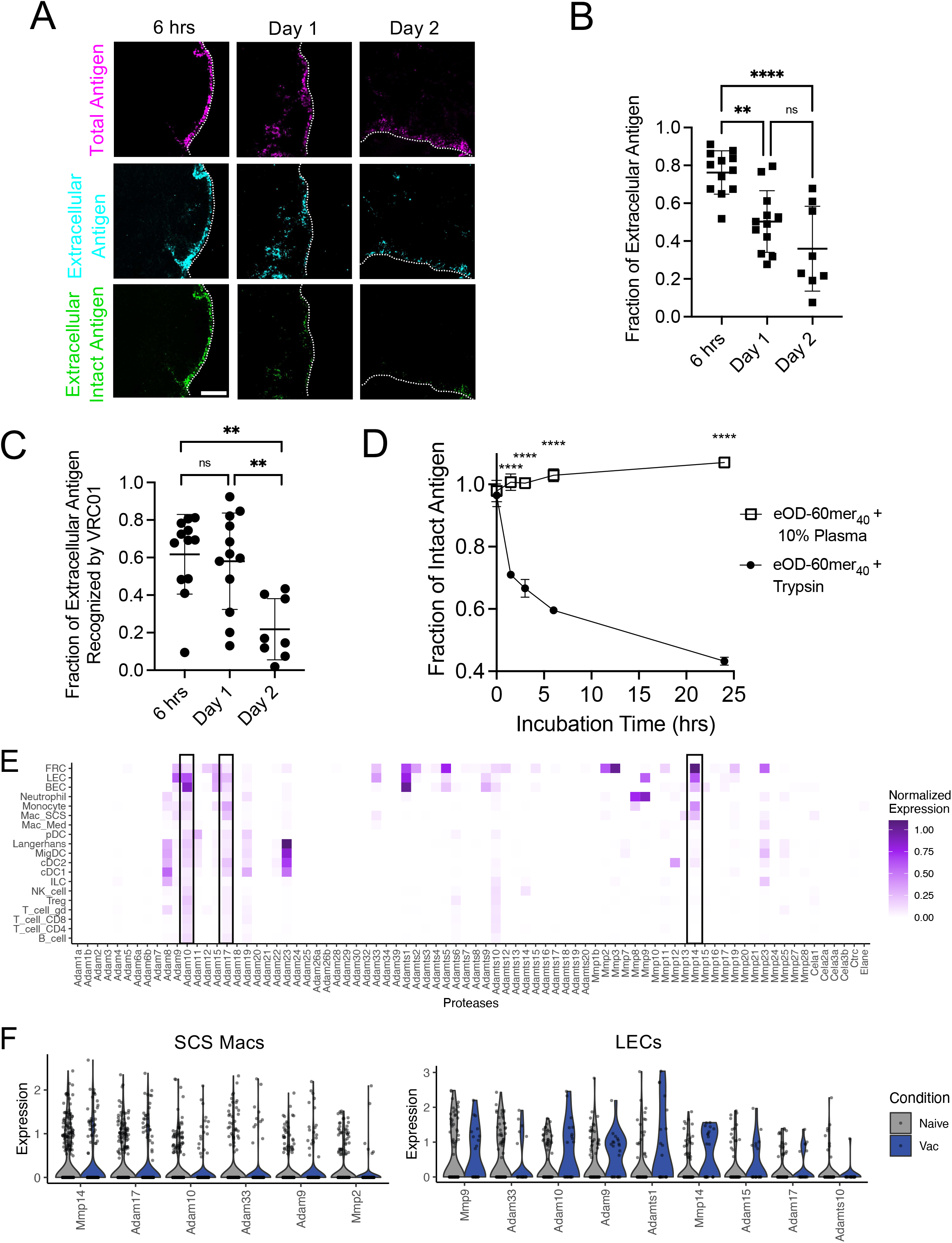
Macrophages and lymphatic endothelial cells lining the lymph node subcapsular sinus express ADAM and MMP proteases. (A-C) C57BL/6J mice (n=3/group) were immunized with 10 µg eOD-60mer_40_ and 5 µg saponin adjuvant and at indicated times LNs were harvested, flash frozen, and sectioned. (A) Shown are representative immunohistochemical images of fixed, non-permeabilized tissue sections showing the total antigen present (top row, detected from the Cy5 fluorescence of the labeled 60mer), extracellular antigen (middle row, detected by staining with an anti-Cy3/Cy5 monoclonal antibody), and intact intracellular antigen (bottom row, detected by VRC01 antibody staining). Scale bar: 100 μm. (B) Fraction of antigen detected in (A) at each time point which was extracellular as determined by colocalization with anti-dye staining. Each point represents one imaged region, collected from at least 6 different tissue sections from 6 independent LNs. Graph shows mean±s.d. (one-way ANOVA with post-hoc Tukey test for multiple pair-wise comparisons, ****p ≤ 0.0001, **p = 0.0017). (C) Fraction of extracellular antigen detected in (A) that was recognized by VRC01 labeling. Each point represents one imaged region, collected from at least 6 different tissue sections from 6 independent LNs. Graph shows mean±s.d. (one-way ANOVA with post-hoc Tukey test for multiple pair-wise comparisons, **p ≤ 0.0032). (D) Plate reader analysis of 10 μg/mL eOD-60mer_40_ incubated with pooled plasma from 2 naïve C57BL/6J mice diluted to 10% v/v in PBS or 10 μg/mL of Trypsin at 37°C for specified amount of time. Time 0 normalized FRET signal (Cy3 excitation and Cy5 emission) divided by Cy5 signal (Cy5 excitation and Cy5 emission) is shown at each time point. Graph shows mean±s.d. (n = 3/group, T-test comparison between 60mer mixed with diluted plasma or Trypsin, ****p ≤ 0.0001). (E-F) C57Bl/6 mice were immunized with 200 µg OVA peptide and 20 µg CpG 1826 adjuvant and LNs were harvested 6 hrs later for analysis by single-cell RNAseq in tandem with LNs recovered from naïve animals. (E) Heatmap of average normalized expression of extracellular protease genes in single cells across lymph node cell types in immunized mice. (F) Normalized expression of the most highly expressed proteases in SCS macrophages and LECs. Arranged in descending order of average expression; protease genes with an average expression value greater than 0.1 are included.

Based on this data, we hypothesized that extracellular proteases expressed by LN cells may play an important role in antigen degradation. To test this idea, we first re-analyzed a single-cell RNA sequencing (scRNA-seq) dataset we recently collected from LNs of naïve or immunized mice, and defined the expression patterns of proteases in murine lymph nodes. We found over 30 genes encoding extracellular or secreted proteases expressed in one or more lymph node cell types (**Figure 2E**). These protease genes were predominantly expressed in stromal cells and myeloid cells and showed low expression in B-or T-lymphocytes. As we observed substantial antigen degradation at the subcapsular sinus, we focused on sinus-lining SCS macrophages and lymphatic endothelial cells (LECs), to identify the most highly expressed proteases in these cell types (**Figure 2F**). The most highly expressed protease genes in SCS macrophages included *Mmp14, Adam17*, and *Adam10*, which encode metalloproteinases; these genes were expressed in both naïve and vaccinated mice. We focused our attention on these 3 proteases as they were also expressed by LECs, and were generally the most broadly/highly expressed across multiple cell types in the LN. *Mmp14* was also highly expressed by fibroblastic reticular cells (FRCs).

To evaluate if expressed protein patterns aligned with the RNAseq profiling, we analyzed the spatial distribution of ADAM17, ADAM10, and MMP14 expression within LNs by immunohistochemistry. All 3 proteases showed high levels of expression along the subcapsular and medullary sinuses as well as within isolated regions in the interior of the LN, and qualitative patterns of expression were the same in resting (**Figure 3A**) and immunized LNs (**Figure S3A-C**). In contrast, the FDC network and its immediate vicinity showed low levels of all 3 proteins (**Figure 3A, S3A-C**). SCS macrophages and LECs showed partial colocalization with all three proteases by immunohistochemistry (**Figure S3D)**. Flow cytometric analysis of permeabilized lymph node cells confirmed the expression of these proteases by LECs and SCS macrophages (**Figure 3B**). Dendritic cells also expressed substantial amounts of ADAM17, but T cells and B cells showed comparatively lower levels of expression of all 3 proteases (**Figure 3B-C**). FDCs, identified by GP38, CR2, and EpCam expression (Duan et al., 2021), also showed little convincing positive staining above the high background observed with isotype control antibody (**Figure 3B)**. We also examined expression patterns of these proteases in human lymphoid tissues. Staining of human tonsil tissue sections showed prominent ridges of ADAM17 and MMP14 expression, with more scattered expression of ADAM10 (**Figure S3E**). However, CD35^+^ follicles were largely devoid of all 3 proteases.

**Figure 3.**
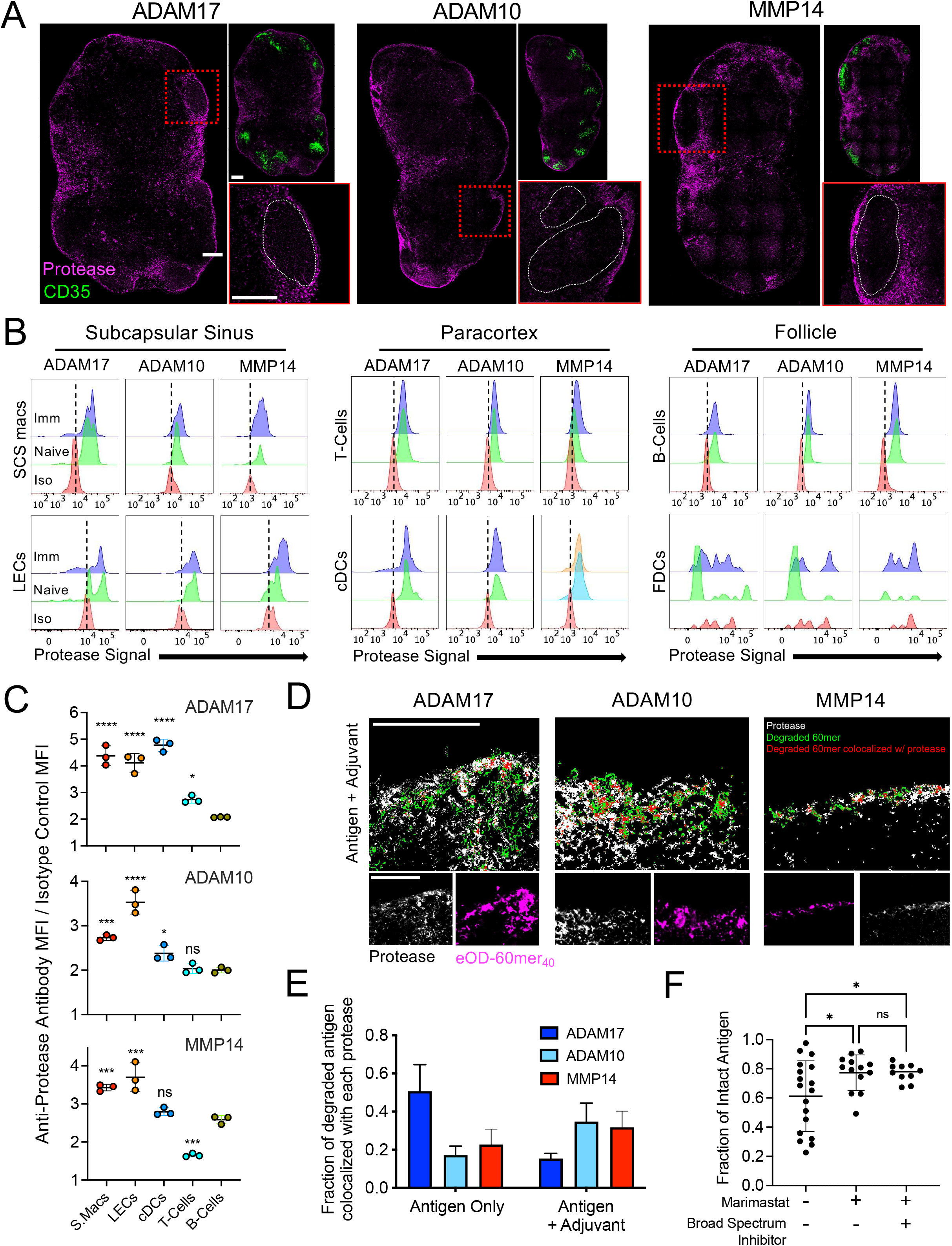
Metalloproteinases are expressed by sinus-lining cells and contribute to rapid antigen degradation in lymph nodes. (A) Resting lymph nodes from C57BL/6J mice (*n*=3 animals/group) were collected for tissue sectioning and stained with monoclonal antibodies to detect metalloproteinases (magenta) as indicated. The bottom right image shows the magnified region within the red dash line along with a white dotted contour denoting B cell follicle. Top right image shows sections with stained protease and the location of the location of the FDC networks (anti-CD35 staining, green). Scale bars: 200 μm. (B-C) LN cells from immunized or naive C57BL/6J mice (*n=*3/group) were isolated, permeabilized, stained with anti-protease antibody, and analyzed by flow cytometry for expression of the indicated metalloproteinases. (B) Representative histograms of ADAM10, ADAM17, and MMP14 expression amongst subcapsular macrophages, LECs, T-cells, DCs, B cells, and FDCs compared to signal from isotype control antibody. (C) The ratio of the MFI of anti-protease to isotype control antibody for each cell type. Shown are mean±s.d. (one-way ANOVA with post-hoc Dunnett test for pair-wise comparison against B cells for multiple pair-wise comparisons, ****p ≤ 0.0001, ***p ≤ 0.001, *p ≤ 0.05). (D-E) C57BL/6J mice (n=3/group) were injected with 10 µg eOD-60mer_40_ for 6 hours or together with saponin adjuvant for 24 hours prior to harvesting the LNs for sectioning and immunohistochemical staining to detect indicated proteases. (D) Regions within slices of LNs injected with saponin adjuvant and antigen for 24 hours showing false color overlays of protease (white), degraded antigen (green) and degraded antigen that colocalizes with protease (red). Degradation of eOD-60mer_40_ was determined by FRET. Scale bars: 100 μm. (E) The fraction of degraded antigen from total detected eOD-60mer_40_ that colocalized with indicated proteases in LNs injected with antigen only or with adjuvant. Data collected from at least 6 tissue sections from 6 lymph nodes. Graph shows mean±s.d. (F) C57BL/6J mice (n=3/group) were immunized with 10 µg eOD-60mer_40_ and 5 µg saponin adjuvant and 2 hours later LNs were vibratome sectioned into 250 µm thick live tissue slices. These tissues were incubated for 6 hours with metalloproteinase inhibitor alone or together with broad spectrum protease inhibitor or left untreated (control). Each point represents one region from one tissue section. Data collected from at least 10 tissue sections from 6 lymph nodes. Graph shows mean±s.d. (one-way ANOVA with post-hoc Tukey test for multiple pair-wise comparisons, *p ≤ 0.028, ****p ≤ 0.0001).

To assess the potential role of metalloproteinases in proteolysis of vaccine antigens, we next assessed whether degrading antigen spatially colocalized with any of these proteases. Tissue sections of LNs from mice injected with only eOD-60mer_40_ 6 hours prior were fixed and stained for CD169 and LYVE-1 to identify SCS macrophages and LECs, respectively. Injected eOD-60mer_40_ co-localized predominantly with macrophages followed by LECs and other cell types not identified by these two markers (**Figure S4A-B)**. To evaluate if metalloproteinases could be contributing to this antigen proteolysis, we examined the colocalization of FRET-labeled eOD-60mer with ADAM17, ADAM10, and MMP14 in the SCS. Mice were immunized with eOD-60mer_40_ alone or together with saponin adjuvant, and using FRET analysis we found that 24 hr later, 20-50% of degraded antigen colocalized with one of these proteases histologically (**Figure 3D-E, S4C**).

To functionally test the role of extracellular protease activity in antigen breakdown, we prepared live vibratome sections of LNs isolated from mice 2 hrs following immunization with eOD-60mer_40_, a timepoint when the majority of antigen that has reached the LN is still intact. These tissue slices were then incubated in the presence or absence of protease inhibitors. In LN slices cultured without drug addition, eOD-60mer degradation proceeded, with on average ∼60% of antigen intact relative to the starting material after 6 hr. Culture of tissue slices with the metalloproteinase-specific inhibitor Marimastat reduced this degradation by ∼50% (**Figure 3F**). Moreover, media containing Marimastat supplemented with a broad spectrum protease inhibitor cocktail against serine, cysteine, aspartic acid proteases as well as amino-peptidases did not yield significantly more intact antigen (**Figure 3F)**. Together these data suggest proteases, and particularly metalloproteinases, contribute to rapid extrafollicular antigen breakdown in lymph nodes.

### Extracellular protease activity is enriched in sinus-lining macrophages and stromal cells but low within follicles

We next sought to understand why antigen localized to FDCs was protected from proteolysis. We observed low staining of ADAM17, ADAM10, and MMP14 in follicles, but this data could not rule out expression of other proteases in the vicinity of FDCs. In addition, although we detected low levels of metalloproteinase expression in permeabilized B cells by flow cytometry (**Figure 3B**), these proteases may not be relevant if they remain intracellular. Thus, to gain a more complete understanding of the spatial pattern of extracellular protease activity in LNs, we employed an imaging zymography approach to visualize protease activity more comprehensively on live tissue sections (**Figure 4A**) (Soleimany et al., 2021a). Live LN vibratome sections were incubated with two fluorescent peptide probes: The first, an activatable zymography probe (AZP), is comprised of a 5-carboxyfluorescein dye linked to a cationic polyarginine (polyR) sequence, followed by a broad-spectrum MMP-cleavable protease substrate sequence and an anionic polyglutamic acid (polyE) sequence. The AZP initially has its cationic polyR arm complexed with the polyE arm, but when proteases cleave the target site of the AZP, the polyR sequence is freed and electrostatically binds to nearby cells, labeling the site of protease activity (**Figure 4A**). The AZP probe was verified to be cleaved by metalloproteinases including ADAM17, ADAM10, and MMP14, as well as aspartic and cysteine protease cathepsins reported to be extracellularly active and expressed by immune cells (**Figure S4D**) (Jakos et al., 2019; Soleimany et al., 2021a; Yadati et al., 2020). Hence, this AZP reports on a broad range of proteases expressed by immune or stromal cells in lymphoid tissues. A second control probe, a Cy5-conjugated polyR peptide, provides a map of overall electrostatic binding sites in the tissue. Tissue areas labeled with both probes identify proteolytically active sites, while locations labeled by the control peptide but not the AZP indicate areas devoid of protease activity.

**Figure 4.**
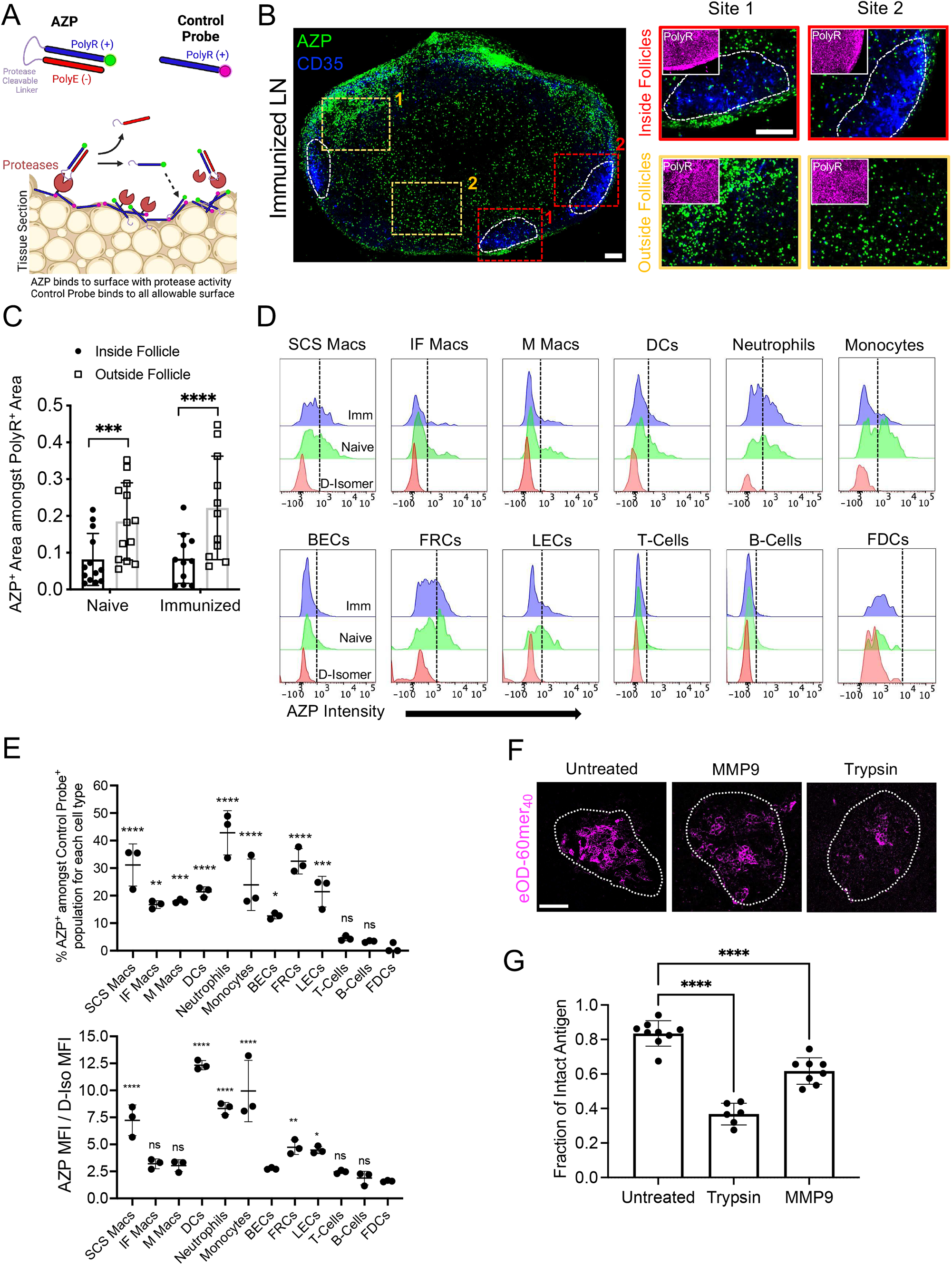
Protease activity is spatially variant in lymph nodes, with high levels in the subcapsular sinus and low activity in B cell follicles. (A) Schematic of fluorescent imaging-based spatial zymography method using activatable zymography probes (AZPs) and control probes to detect protease activity on live tissue sections. (B-E) C57BL/6J mice (n=3/group) were left naive or immunized with 10 µg eOD-60mer and 5 µg saponin adjuvant. LNs were harvested 24 hr later and vibratome sectioned into 250 µm thick live tissue slices that were incubated with AZP and control probe for 2 hours, followed by overnight fixation for confocal imaging (B-C) or processing for flow cytometry analysis of probe binding (D-E). (B) Representative whole-LN tissue section (left) and higher-magnification images (right) taken from boxed regions around FDC networks (“Inside Follicle”) or in extrafollicular regions (“Outside Follicle”). Main images show AZP probe binding (green) overlaid on anti-CD35 staining (blue), while insets at right show the corresponding staining from the same areas for the control PolyR probes (magenta). Regions encircles by white dashed lines indicate follicle regions used for analysis. (C) Quantification of relative AZP probe binding “inside follicles” vs “outside follicles”. Each point represents one LN section, collected from a total of 11-13 LN sections from 6 independent LNs. Graph shows mean±s.d. (T-test comparison between AZP binding “inside follicles” and “outside follicles”, ***p ≤ 0.001, ****p ≤ 0.0001). (D) Representative flow cytometry histograms for AZP binding to immunized (Imm) or resting LN (Naïve) cells, and D-isomer control probe binding to resting LN cells. Dashed line provides a guide to the eye denoting AZP positive (AZP^+^) and negative (AZP^-^) populations. (E) Quantification of AZP probe binding (top graph) and ratio of AZP to D-Isomer MFI (bottom graph) amongst LN cell populations as determined by flow cytometry. Graph shows mean±s.d. (one-way ANOVA with post-hoc Dunnett test for pair-wise comparison against AZP^+^ FDC percentage, ***p ≤ 0.01, ***p ≤ 0.001, ****p ≤ 0.0001). (F-G) C57BL/6J mice (n=3/group) were immunized as in (B-E) and LNs were harvested 3 days later. LNs were vibratome sectioned into 250 µm thick live tissue slices, which were incubated for 4 hours with trypsin, MMP9, or left untreated. (F) Representative images of follicles post treatment showing eOD-60mer remaining on the FDC network. Encircled region within dashed white lines indicates the boundary of the B cell follicle. (G) Quantification of FDC-localized antigen detected on untreated, trypsin-treated, or MMP9-treated LN sections. Each point represents one region from one tissue section. Data collected from at least 6 tissue sections from 6 lymph nodes. Graph shows mean±s.d. (one-way ANOVA with post-hoc Dunnett test for pair-wise comparison against Untreated, ****p ≤ 0.0001). All scale bars indicate 100 μm in length.

LN tissue sections from naïve animals or mice immunized with eOD-60mer and adjuvant showed AZP labeling of discrete cells across the entire tissue–with particular concentrations of protease activity at the SCS–except within the immediate vicinity of the FDC networks (**Figure 4B, Figure S4E**). This was not due to a lack of binding sites for the AZP in follicles, as the control polyR peptide bound at high levels across these regions (**Figure 4B, S4E** insets). Incubation of LN sections with a control uncleavable D-Isomer form of the same AZP peptide sequence showed substantially lower fluorescence signal, suggesting that AZPs were not internalized and required extracellular proteolysis to bind to cells (**Figure S4F-H**). Quantification of AZP signal within FDC regions vs. adjacent LN tissue revealed substantially lower protease activity in the follicles, and this pattern of protease activity distribution was qualitatively similar for resting and immunized lymph nodes (**Figure 4C**).

We next sought to phenotypically identify cells with active protease activity defined by AZP labeling. Flow cytometry analysis of single cells extracted from PolyR- and AZP-treated LN sections revealed that 80% or more of all lymphocyte and stromal cell populations examined were positive for control PolyR binding (**Figure S5A-C**) (Soleimany et al., 2021b). Gating on cells positive for PolyR binding (indicating sufficient binding sites for AZP-based protease detection), we then evaluated the relative levels of protease activity as detected by AZP binding. Neutrophils, SCS macrophages, and fibroblastic reticular cells (FRCs) had the highest levels of AZP binding above the background labeling by D-isomer control AZP in both resting (**Figure S5D**) and immunized LNs (**Figure 4D-E)**. All macrophage subsets examined (CD169^+^CD11b^hi^F4/80^low^ subcapsular; CD169^+^CD11b^low^F4/80^low^ interfollicular; and CD169^+^F4/80^hi^ medullary macrophages) had some proportion of AZP^+^ cells, but SCS macrophages showed the most prominent protease activity (**Figure 4D-E, Figure S5D**). Lymphatic endothelial cells (LECs), DCs, and monocytes also exhibited lower but clearly detectable protease activity, while FDCs, B cells, and T cells showed very low levels of AZP labeling **(Figure 4D-E, Figure S5D)**. Taken together, these results suggested that many sinus-lining cells have high levels of protease activity, while cells in the follicle exhibit low levels of extracellular proteolysis.

Although the spatial zymography assay suggested protease activity in follicles is substantially reduced relative to extrafollicular regions of the LN, we also assessed additional potential mechanisms of antigen protection within follicles. As eOD-60mer is recognized by MBL and becomes complement-decorated in the presence of serum, it was possible that MBL and/or complement binding might have a shielding effect blocking proteolytic attack on FDC-localized antigen. However, in an *in vitro* trypsin digestion assay, eOD-60mer was proteolyzed with destruction of the VRC01 epitope regardless of serum-derived C3 binding on the NP surface, suggesting that MBL or complement binding to the particles cannot protect the antigen from protease activity (**Figure S5E**). A second possibility could be that antibodies produced very early following immunization form immune complexes with the antigen on FDCs to sterically inhibit protease attack, or that FDCs capturing the antigen rapidly internalize and recycle it on their dendrites, providing a degree of protection from protease exposure in the follicles. We immunized MD4 mice where B cells transgenically express an antigen receptor specific for an irrelevant antigen with eOD-60mer_40_, and found antigen localized to FDCs after 2 days was largely non-degraded similar to wild type mice, suggesting that antibody-mediated protection of antigen is not a factor at least at these early time points (**Figure S5F**). To assess whether FDCs could protect antigen in the face of artificially enforced protease activity in follicles, we immunized mice with eOD-60mer_40_ and adjuvant, extracted lymph nodes 3 days later when antigen was concentrated on the FDC networks, and incubated live LN slices with trypsin or recombinant matrix metalloproteinase (MMP9) to expose FDC-localized antigen to protease attack. Both trypsin and MMP9 treatment led to significant loss of intact eOD-60mer within 4 hrs, suggesting that antigen localized to FDCs is not intrinsically protease resistant (**Figure 4F-G**). Hence, while additional factors may play a role, a lack of active protease action in the vicinity of FDCs appears to be an important contributor to the long lifetime of immunogens captured in the FDC networks.

### Immunization regimens promoting antigen localization to FDCs amplify B cell responses to intact antigen epitopes without increasing B cell targeting of antigen breakdown products

Rapid antigen degradation in extrafollicular regions of the LN could expose non-native or irrelevant internal epitopes of immunogens to B cells and thereby limit responses to the intact antigen. Conversely, we hypothesized that immunization regimens promoting rapid delivery of antigen to FDCs might enhance responses to the native immunogen. To test this idea, we carried out experiments using a stabilized HIV Env gp140 SOSIP trimer termed N332-GT2 in soluble trimer and protein nanoparticle forms (Steichen et al., 2019). We first analyzed the degradation behavior of trimers using our FRET methodology. Labeling conditions were identified (6 dyes per trimer on average, Trimer_6_) that allowed for FRET tracking of N332-GT2 integrity with minimal perturbation of the antigenicity profile of the trimer (**Figure S6A**). Similar to our findings with eOD-60mer, the FRET efficiency of Trimer_6_ was highly correlated with the proportion of intact antigen as determined by PAGE analysis following trypsin digestion *in vitro* (**Figure S6B-D)**. Loss of FRET signal on protease-treated trimer also correlated with reduced binding of structure-sensitive antibodies, especially the bnAb 35O22 that recognizes the gp120-gp41 interface (**Figure S6E**). Concomitantly, trypsin-digested Trimer_6_ began to show binding of antibodies that recognize epitopes of the gp120 inner domain that are buried in the interior of the properly folded trimer (**Figure S6E**). Similar to our findings with eOD-60mer, FRET-labeled N332-GT2 trimer showed no signs of degradation when exposed to serum for at least 24 hrs (**Figure S6F**). In addition to the soluble trimer format, this trimer can be produced as a NP by fusion of the trimer sequence with ferritin subunits (abbreviated Trimer-NPs) (Steichen et al., 2019). For N332-GT2 nanoparticles, 40 dyes could be conjugated to each particle (Trimer-NP_40_) without impacting binding of trimer structure-specific mAbs (**Figure S6G**).

Using these FRET-labeled immunogens, we examined the localization and *in vivo* stability of trimers administered under different dosing regimens (**Figure 5A**). We previously showed that ferritin-based trimer-NPs are rapidly transported to FDCs via the same MBL/complement pathway as the eOD-60mer immunogen, while soluble trimer distributes more diffusely in LNs prior to clearance (Martin et al., 2021; Tokatlian et al., 2019). Consistent with those prior findings, soluble Trimer_6_ or Trimer-NP_40_ delivered as bolus injections were detected in the SCS for up to two days, but Trimer-NP_40_ also exhibited accumulation on FDCs starting 2 days post immunization (**Figure 5B-D**). We also administered soluble Trimer_6_ using an “escalating-dose” (ED) immunization (Cirelli et al., 2020; Tam et al., 2016) administering the same total antigen/adjuvant dose as the other two immunizations, but administered as 7 injections of increasing dose over two weeks (**Figure 5A**). ED immunization is another approach that promotes antigen deposition on FDCs, through immune complex formation as the dosing progresses (Cirelli et al., 2020; Tam et al., 2016), and we found that 2 days after the last injection (day 14) of Trimer_6_ by ED, substantial amounts of the antigen were localized on the FDC network (**Figure 5B-D**). Like eOD-60mer, FRET analysis of Trimer_6_ and Trimer-NP_40_ indicated that substantial portions of extrafollicular antigen (50% of Trimer and 80% Trimer-NP) were degraded within 2 days, while Trimer-NP_40_ or ED-administered Trimer_6_ localized to FDCs remained predominately intact (**Figure 5E**). Thus, immunization with HIV Env trimers using extended-dosing regimens or nanoparticle forms promotes follicular targeting and increases the level of intact antigen retained within the LN.

**Figure 5:**
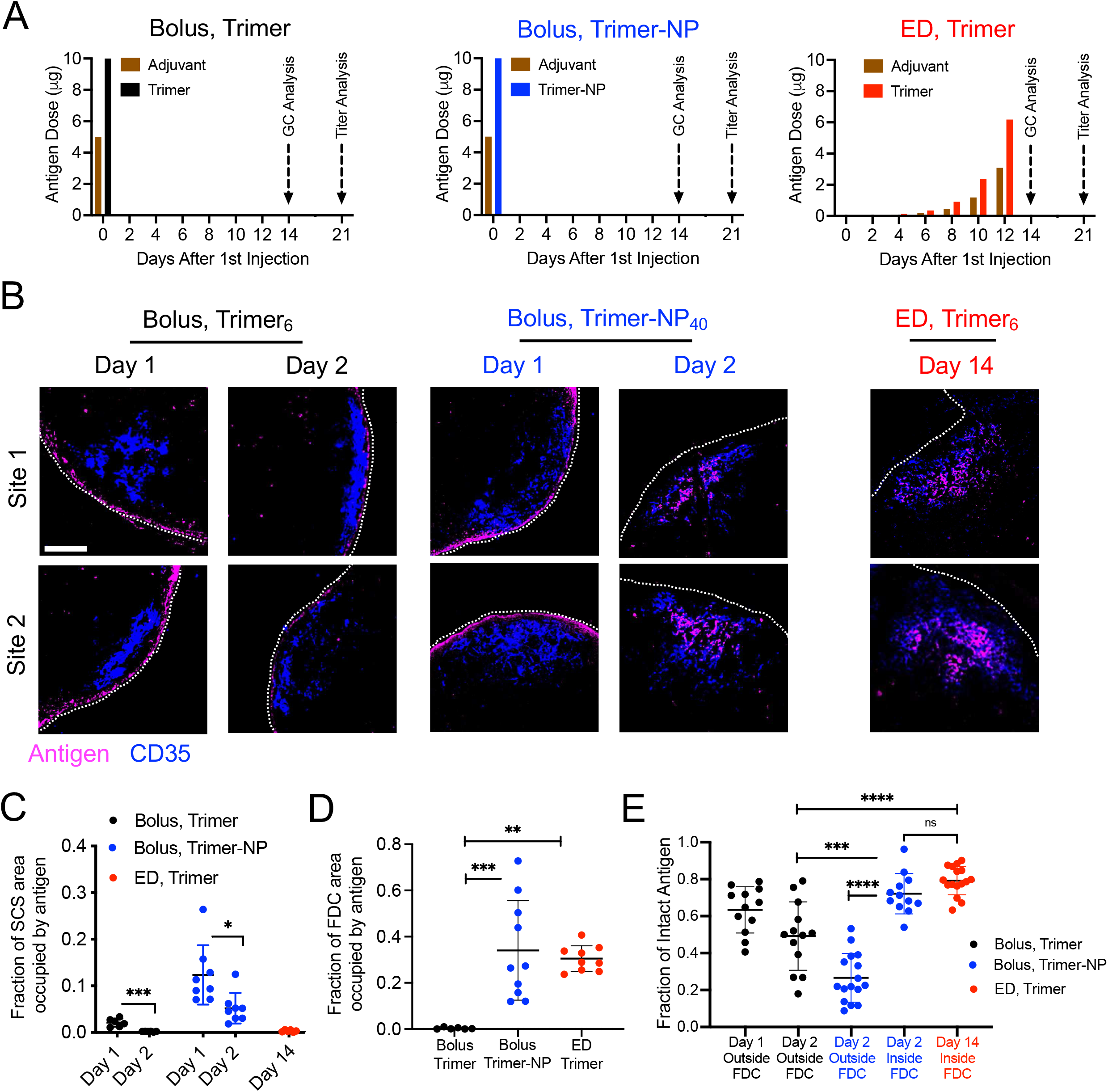
Immunizations targeting antigen to FDCs preserve antigen integrity *in vivo*. (A) Administration schedules for bolus and escalating dose (ED) immunization regimens using soluble Trimer (Trimer) and Trimer-ferritin (NP) immunogens. Saponin adjuvant was used in all conditions. (B-E) C57BL/6J mice (n=3/group) were immunized with indicated antigens and dosing scheme. At indicated time points, LNs were harvested, flash frozen, and sectioned for confocal imaging, antigen localization, and FRET analysis. (B) Representative confocal images of follicular regions within lymph node sections with dye labeled antigens (magenta) and anti-CD35 stain (blue) for FDC networks. Two distinct representative locations are shown for each immunization condition; white dotted line indicates the edge of the tissue section. Scale bar: 100 μm. (C) Quantification of antigen localized to the subcapsular sinus at indicated times. Each point represents one tissue slice. Data collected from at least 6 LNs. Graph shows mean±s.d. (T-test comparison between Day 1 and Day 2 for each immunization condition, ***p = 0.0003, *p = 0.0136). (D) Fraction of antigen localized to the FDCs at day 2 for Bolus Trimer and Bolus NP immunization and day 14 (two days after last dosing) for ED Trimer. Each point represents one tissue slice. Data collected from at least 6 lymph nodes. Graph shows mean±s.d. (one-way ANOVA with post-hoc Tukey test for multiple pair-wise comparisons, ***p = 0.0004, **p = 0.0015). (E) Mean fraction of intact labeled antigens in the lymph node at the indicated locations and times as determined by FRET imaging. Each point represents one region from one tissue section. Data collected from at least 8 tissue sections from 8 lymph nodes. Graph shows mean±s.d. (one-way ANOVA with post-hoc Tukey test for multiple pair-wise comparisons, ****p ≤ 0.0001, ***p = 0.0003).

We next sought to determine the impact of FDC localization and enhanced antigen integrity on B cell responses by tracking germinal center responses to intact trimer vs. off-target breakdown products. Bolus immunization with Trimer-NP or ED immunization with Trimer led to modestly increased 2- and 3-fold greater total GC B cell frequencies compared to bolus free trimer immunization, respectively (**Figure 6A, B**). However, when GC B cells were stained with N332-GT2 trimer probes to identify cells capable of binding to the intact antigen at day 14, much larger differences were observed, with ∼30% of GC B cells specific for the intact trimer following ED immunization and ∼10% trimer-specific GC B cells following trimer-NP immunization, while only ∼2% of GC cells bound to intact trimer following bolus N332-GT2 trimer vaccination (**Figure 6B**). To ensure the accuracy of this measurement, trimer probes were also used to stain GC B cells from mice immunized with an irrelevant antigen, ovalbumin (Ova), and this control showed low background staining (**Figure 6B**). Longitudinal analysis showed that the proportion of trimer-specific GC B cells following bolus trimer vaccination remained relatively constant through 21 days post immunization and never reached the levels primed by Trimer-NP or ED vaccination (**Figure 6C**).

**Figure 6:**
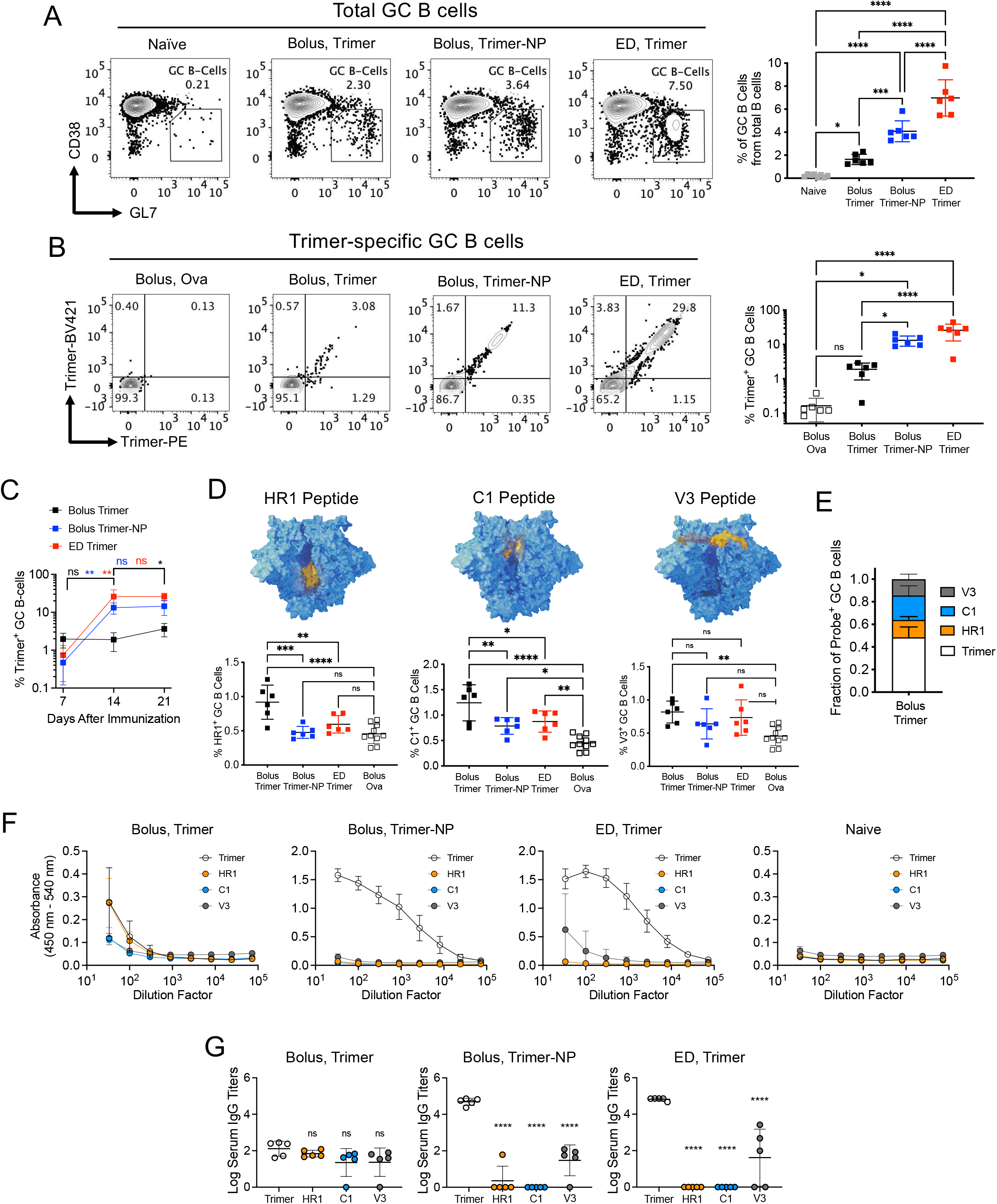
Targeting of immunogens to FDCs amplifies GC B cell responses against intact antigens without increasing the response to antigen breakdown products. C57BL/6J mice (n*=*5-8/group) were immunized with indicated antigens and dosing schemes as defined in Figure 5A. (A-D) LNs were harvested at day 14 and processed for flow cytometry analysis of GC B cells. (A) Representative flow cytometry plots showing proportions of GC B cell populations (left) and percent of GC B cells from total B cells under different immunization conditions (right). Graph shows mean±s.d. (One-way ANOVA with post-hoc Tukey test for pair-wise comparison between all groups, ****p ≤ 0.0001, ***p ≤ 0.001, *p ≤ 0.05). (B) Representative flow cytometry plots showing Trimer-specific GC B cells (left) and percent of Trimer-specific GC B cells under different immunization conditions (right). Graph shows mean±s.d. (One-way ANOVA with post-hoc Tukey test for pair-wise comparison between all groups, ****p ≤ 0.0001, *p ≤ 0.05, ns = not significant). (C) Time course analysis of trimer-specific GC B cells for indicated immunization condition. Graphs show mean±s.d. (One-way ANOVA with post-hoc Tukey test for pair-wise comparison between different time points for each immunization condition, **p ≤ 0.01, *p ≤ 0.05, ns = not significant). (D) Structural models of the HIV Env trimer immunogens highlighting (in yellow) the location of the HR1 peptide, C1 peptide, and V3 peptide domains used as probes to identify breakdown product-specific GC B cell populations and antibody responses (top) and percent of GC B cells specific for the indicated probes on day 14 post immunization (bottom). GC B cells from Bolus Ova immunized mice stained with different probes showed similar background staining and are shown as an aggregate in these plots. Graph shows mean±s.d. (One-way ANOVA with post-hoc Tukey test for pair-wise comparison between Bolus Ova, Bolus Trimer, Bolus Trimer-NP, and ED Trimer, ***p ≤ 0.001, **p ≤ 0.01, *p ≤ 0.05, ns = not significant). (E) Quantification of the fraction of GC B cells specific for each of the 4 antigen probes for Bolus Trimer immunization regimen. Graph shows mean±s.d. (F) Raw mean ELISA absorbance vs. serum dilution (starting at 3% serum in blocking buffer) for each immunization regimen showing differential binding to intact trimer and off-target probes. Serum was collected 21 days after initial dose. Graphs show mean±s.d. (G) Endpoint serum antibody titers against trimer and off-target probes 21 days after initial dose. Shown are mean±s.d. (One-way ANOVA with post-hoc Dunnett test for pair-wise comparison of the off-target probes to intact trimer, ***p < 0.001, ****p < 0.0001, ns = not significant).

This highly amplified response against the intact immunogen using NP or ED immunization could reflect the fact that GC B cells are encountering antigen in a multivalent form (either the NP, or as immune complexes containing many trimers in the case of ED immunization). However, we also investigated the frequency of GC B cells reacting to trimer breakdown products, by staining GC B cells with HR1, C1, or V3 loop peptide probes, spanning different regions of the gp140 protomers that are buried in the intact trimer (HR1, C1, V3) (**Figure 6D, S6H)**. To define the detection limits of this assay, GC B cells from mice immunized with the irrelevant antigen ovalbumin (Ova) were also analyzed with these probes. Fourteen days post immunization, low but statistically significant levels of GC B cells recognizing the C1 epitope were detectable following NP and ED immunizations, but responses to the HR1 or V3 peptides were not detected above the Ova background control (**Figure 6D, S6H**). By contrast, bolus trimer immunization led to GC responses targeting each of these internal epitopes that were similar in magnitude to the total response to intact trimer (**Figure 6B, 6D, S6H**), and in fact the sum of responses to these irrelevant epitopes was slightly greater than the total GC B cell response recognizing intact trimer (**Figure 6E**). Thus, although multivalency of the antigen achieved by NP or ED immunization potentially plays an important role in amplifying the trimer-specific GC response, it does not amplify breakdown product-specific responses in tandem, consistent with the idea that localization to FDCs preserves antigen integrity.

We also examined whether the trends observed in GC responses were reflected in subsequent serum antibody titers. Compared to bolus trimer immunizations, NP or ED yielded 1000-fold higher serum IgG titers recognizing the intact trimer 3 weeks post immunization (**Figure 6F-G**). Further, the low titers against intact trimer observed after bolus trimer immunization were of the same order of magnitude as responses to the breakdown product epitopes from HR1, C1, and V3 peptide. By contrast, titers against the internal epitope peptides were undetectable following NP or ED vaccination, with the exception that ED immunization resulted in low responses against the V3 peptide (**Figure 6F-G**). These results collectively illustrate that by designing immunization regimens to favor antigen localization to FDCs, GC and antibody responses to breakdown products can be reduced, while responses against structurally-intact antigen can be greatly amplified.

## DISCUSSION

Elicitation of protective antibody responses by vaccination is predicated on the activation and affinity maturation of B cells whose antigen receptors recognize epitopes present on the functional target protein of the targeted pathogen. Toward this goal, many elegant advances in protein engineering in recent years have led to the generation of vaccine immunogens that present neutralizing epitopes faithfully mimicking the structure of viral native envelope proteins and exhibiting greatly enhanced structural stability relative to the native antigens (Graham et al., 2019; Ng’uni et al., 2020). However, despite the critical importance of immunogen structure to the specificity of the resulting immune response, the lifetime of structurally-intact antigen *in vivo* following immunization is poorly understood. Here we applied a FRET imaging-based approach to track the fate of several types of clinically relevant protein immunogens in mice, leading to several important findings: First, antigens arriving in the subcapsular sinus and extrafollicular regions of lymph nodes rapidly begin degrading, with significant losses in integrity within 24-48 hr. We found that this occurred even for highly stabilized HIV Env trimers and was independent of protease activity in serum. Second, by contrast, immunogens that localize to the dendrites of FDCs within this early time window remain largely intact and are preserved for more than a week at these sites. We characterized lymph node protease activity and protease expression at the transcriptional and protein levels, and discovered spatial heterogeneity in protease activity, with high protease levels and activity found in stromal cells and macrophages, especially in the sinuses, but low levels of protease activity in B cell follicles around the FDC networks. Based on these findings, we modulated the delivery of antigens to FDCs by using different immunogen formats or vaccination approaches, and found that traditional bolus immunization using soluble HIV envelope antigens led to weak GC B cell responses to the structurally-intact immunogen, which were matched in magnitude by responses against irrelevant breakdown products. However, using immunizations that promoted antigen accumulation on FDCs, responses to intact antigen could be greatly amplified without concomitant increases in responses to off-target antigen breakdown products.

While protease activity in LNs is understudied, antigen degradation within the SCS has been reported. Jenkins and colleagues observed that immunization with fluorescent hen egg lysozyme (HEL) conjugated to polymer microspheres labeled in a second color led to significant antigen acquisition by HEL-specific B cells within 4 hours of injection in the absence of microsphere uptake, suggesting rapid proteolytic release of antigen from the particles *in vivo* (Catron et al., 2010). *In vitro* serum exposure experiments suggested HEL could be cleaved from the microparticles by plasma proteases. In the present experiments, we detected intact HIV eOD and Env trimer nanoparticles within the SCS well over 4 hours after immunization, and incubation of eOD-60mer or the stabilized Env trimer with diluted plasma resulted in minimal degradation of the antigen as determined by FRET analysis. Susceptibility to plasma proteases may be antigen-specific, and immunogen design efforts to stabilize these antigens and/or the dense glycosylation on HIV envelope proteins may provide these immunogens with added protection (Wagh et al., 2020).

Antigen retention and protection within FDCs was first suggested over three decades ago in seminal work by Tew and colleagues. Immunogen stability was shown indirectly through chromatography after retrieval of radiolabeled antigen from whole LNs (Tew et al., 1979). However, with the emerging evidence that lymphatic endothelial cells can also store antigen in a complement receptor 2-independent manner, antigen retrieved from whole lymph node extracts may not be specific to FDCs (Tamburini et al., 2014). Monoclonal antibodies have been used to detect antigens within FDCs (Bachmann et al., 1996), but this approach can only report on individual epitopes and is limited by the potential for vaccine-elicited antibody response to confound interpretation. Experiments such as these have led to the speculation that the FDC network may provide a sanctuary site that not only retains antigen but protects it from degradation (Cyster, 2010), but direct evidence in support of this idea has remained sparse. Using FRET analysis to report on overall antigen integrity, we find that immunogens localized to FDCs during the early stages of the humoral response retain near constant levels of integrity over at least 7 days *in vivo*. Consistent with these findings, AZP-based spatial zymography assays applied to live tissue sections suggested that extracellular protease activity is low in follicles relative to the SCS and interfollicular regions of lymph nodes. Prior work has shown that FDCs cyclically present and internalize captured antigens from their surfaces, which would be expected to aid in protecting antigen from proteolytic activity locally (Heesters et al., 2013), and recently published *in silico* modeling suggests that the kinetics of antigen recycling is crucial for preserving antigens in GCs (Arulraj et al., 2021; Heesters et al., 2013). However, when we challenged LN sections with recombinant proteases, FDC-localized antigen underwent degradation, indicating that this recycling process would be inefficient for protecting antigen in the face of significant local protease activity in the follicle. Nevertheless, it seems likely that antigen recycling and low protease activity in follicles work together to preserve antigen on FDCs over prolonged periods.

Outside of B cell follicles, transcriptional analysis, immunohistochemistry, and flow cytometry identified substantial expression of ADAM and MMP proteases, particularly on cells lining the subcapsular sinus. ADAMs and MMPs are well known for their sheddase activity and ECM degradation, respectively, but studies suggest that their substrate list extends well beyond these functions. ADAM17 has been reported to cleave Ebola virus glycoproteins and soluble fibronectin, while MMP14 sheds cell surface receptors such as ICAM, CD44 and cytokines such as IL-8 (Calligaris et al., 2021; Itoh, 2015). These observations suggest that metalloproteinases are promiscuous with their choice for substrates, and this is reflected in their consensus sequence that is highly variable except for the P1 and P1’ amino acid locations that flank the cleavage site (Scharfenberg et al., 2020). Based on mass spectrometry analysis, these amino acids are identified to be AV or AL for both ADAM10 and ADAM17 (Scharfenberg et al., 2020) while MMP14 and MMP9 both recognize GL (Rawlings et al., 2014). For the immunogens tested here, these motifs are present in the peptide sequences of the outer eOD and inner lumazine synthase core of the 60mer nanoparticle and are scattered across the HIV Env trimer sequences. In addition, ADAMs often cleave cell surface-bound proteins, with some reports of activity on soluble substrates, while MMP14 can cleave proteins proximal and distal to the cell membrane (Itoh, 2015). Hence, these proteases seem likely to be capable of degrading antigens generally.

Recent studies have begun to interrogate the antigen specificity of germinal center B cells during ongoing GC responses following immunization, and have shown that often, a minority of GC B cells can be shown to have affinity for the intact immunizing immunogen through staining with fluorescent multimeric probes (Kuraoka et al., 2016; Moyer et al., 2020; Tokatlian et al., 2019). The remaining B cells might be GC cells that have made mutations leading to a non-functional antigen receptor but have not yet committed to apoptosis (Shlomchik et al., 2019), they might have specificities to undefined products (which have been referred to as recognizing “dark antigen” (Kuraoka et al., 2016)) or alternatively, might be antigen-specific but have affinities too low for detection by simple staining methods (Viant et al., 2020). By staining GC B cells with probes representing buried epitopes in intact HIV Env trimers, we demonstrate here that following bolus trimer immunization, which fails to deliver substantial antigen to FDCs, the population of GC B cells responding to breakdown products is as large as the population recognizing the intact immunogen. Such a substantial response against irrelevant epitopes will be problematic for difficult pathogens such as HIV, where B cells capable of maturing to produce broadly neutralizing antibodies are very rare (Jardine et al., 2016), because competition in germinal centers can overwhelm these rare precursors (Abbott et al., 2018; Dosenovic et al., 2018). Contrasting these results, immunization with nanoparticle forms of the immunogen or utilizing “escalating dosing” immunization to promote antigen trafficking to FDCs greatly amplified the intact antigen-specific GC B cell response. Retention of intact antigen by FDCs plays an important role in this outcome because no parallel increase in responses to breakdown products was detected in these immunizations. These data align well with the finding in non-human primate immunization studies that ED regimens increase the induction of autologous tier 2 neutralizing responses to Env trimer immunization (Cirelli et al., 2020).

In summary, the studies presented here provide evidence for rapid proteolysis and clearance of antigens from extrafollicular regions of lymph nodes following immunization, but long-lived retention of structurally-intact antigen by follicular dendritic cells. This data provides clear motivation for immunization strategies that promote antigen delivery to the FDC network. While we demonstrated two approaches to achieve this, via nanoparticle delivery or escalating dosing immunization, a variety of approaches may be capable of achieving this goal, such as the immunization with immune complexes (Wang et al., 2019), use of antigen-complement fusions (Dempsey et al., 1996) or extended antigen delivery via nucleic acid vaccines (Pardi et al., 2015). Such approaches may be particularly important for successful generation of protective antibody responses against difficult pathogens such as HIV.

## ACKNOWLEDGEMENTS

We would like to thank the Koch Institute Swanson Biotechnology Center’s Flow Cytometry and Microscopy core facilities for their technical support. This work was supported in part by the NIH (award UM1AI144462 to W.R.S. and D.J.I.; awards AI125068 and P01AI048240 to D.J.I.), Koch Institute Support Grant P30-CA14051 from the National Cancer Institute (S.N.B.), a Core Center Grant P30-ES002109 from the National Institute of Environmental Health Sciences (S.N.B.), the Ragon Institute of MGH, MIT, and Harvard (D.S.K. and D.J.I.), and the Marble Center for Cancer Nanomedicine (S.N.B. and D.J.I.). A.A. was supported by NIH Training grant, 5T32AI007386. A.P.S. acknowledges support from the NIH Molecular Biophysics Training Grant and the National Science Foundation Graduate Research Fellowship. J.D.K. acknowledges fellowship support from the Ludwig Center at MIT’s Koch Institute. D.J.I. and S.N.B. are Howard Hughes Medical Institute Investigator.

## AUTHOR CONTRIBUTIONS

Conceptualization, A.A. and D.J.I. Methodology, A.A., D.J.I., A.C., N.H., A.P.S., J.D.K., and S.N.B. Software, A.A. and H.L. Formal Analysis, A.A., A.C., and H.L. Investigation, A.A., M.B., A.C., H.L., M.S., J.R.G., P.A., T.R., S.X., W.A., J.A., and H.S. Resources, A.C., N.H., A.P.S., J.D.K., S.N.B., C.A.C., W.R.S., L.M.F., P.H., and D.S.K. Writing – Original Draft, D.J.I., and A.A. Writing – Review & Editing, D.J.I., A.A., W.R.S., N.H., S.N.B., and D.S.K. Visualization, A.A., D.J.I., and H.L. Funding Acquisition, D.J.I.

## MATERIALS AND METHODS

### Mice

Six-nine week old female C57BL/6J mice were purchased from Jackson Laboratory (Strain No: 000664). Tg(IghelMD4)4Ccg/J breeding pairs were purchased from Jackson Laboratory (Stock No: 002595) and mated in-house. Their progeny was genotyped (Transnetyx) and female mice between ages of 8-11 weeks old were used for immunization studies.

### Saponin Adjuvant preparation

The saponin adjuvant used in this study is an ISCOMs-like self-assembled nanoparticle comprised of cholesterol, phospholipid, and Quillaja saponin as previously described (Tokatlian et al., 2019). Briefly, under sterile conditions, the following solutions were made: 0.5 mL of 20 mg/mL of cholesterol (700000P, Avanti) in 20% MEGA-10 detergent (D6277, Sigma Aldrich), 0.5 mL of DPPC (850355C, Avanti) in 20% MEGA-10 detergent, and 0.5 mL of 100 mg/mL Quil-A adjuvant (vac-quil, Invivogen) in deionized H_2_0 were prepared. DPPC solution was mixed with the cholesterol followed by the addition of Quil-A saponin in rapid succession. This mixture was diluted with PBS to a concentration of 1 mg/mL cholesterol and 2% MEGA-10, prior to overnight equilibration at 25°C. The lipids/saponin/surfactant solution was then dialyzed against PBS using a 10 kDa MWCO membrane for 5 days at 25°C and filter sterilized using a 0.2 Supor syringe filter. For further purification, the adjuvant solution was concentrated using 50 kDa MWCO Amicon Ultra-filters (UFC905008, Millipore Sigma) and purified by size exclusion chromatography using a Sephacryl S-500 HR size exclusion column. For quality control, the final saponin adjuvant was characterized by Limus Amebocyte Lystae assay (QCL-1000, Lonza) for low endotoxin levels. The adjuvant concentration was determined using a cholesterol quantification kit (MAK043, Sigma).

### Recombinant immunogen production

eOD-60mer was produced recombinantly as previously described. The eOD-60mer gene comprised of eOD-GT8 monomer fused to Lumazine Synthase was synthesized by Integrated DNA Technologies, cloned into phLsec plasmid, and transfected into Expi293F (A14527, Thermo Fisher Scientific) cells. After 6 days cell culture supernatant was harvested by centrifugation and filtration through a 0.2 μm filter. The eOD-60mer was affinity purified by incubating with *Galanthus Nivalis* Lectin-conjugated agarose beads (AL-1243-5, Vector Laboratories) overnight under gentle agitation at 4°C and eluted with lectin elution buffer containing 1 M Methyl a-D-mannopyranoside (M6882, Millipore Sigma). The resulting solution was dialyzed in PBS and further purified by size exclusion chromatography using Sephacryl S-500 HR resin.

N332-GT2 trimers were expressed in FreeStyle 293F cells (Invitrogen, Cat no. R79007) and purified in two steps by affinity chromatography using a GE HisTrap column and size-exclusion chromatography using a GE S200 Increase column as described previously (Steichen et al., 2016; Steichen et al., 2019). N332-GT2 nanoparticles were expressed and purified as previously described (Steichen et al., 2019).

### Antigen labeling and characterization

Protein antigens (eOD-60mer, HIV Env trimer, or trimer-NPs) at 1 mg/mL in PBS were diluted at a 1:1 volume ratio in 0.2 M Sodium Bicarbonate buffer (S8875, Sigma-Aldrich) pH 8.4 and kept on ice. Fresh 1 mg/mL stock solutions of Sulfo-Cyanine 3 (21320, Lumiprobe) and -Cyanine 5 (23320, Lumiprobe) NHS esters were made in 0.2 M Sodium Bicarbonate (pH 8.4) and added to the antigen solutions. This reaction was allowed to proceed for 16 hr at 4°C then samples were desalted by passing through a Zeba Spin Desalting column (40 kDa Mw Cutoff) equilibrated in PBS twice. Labeled antigens were sterilized by filtering through 0.22 µm pore size Spin-X centrifuge tube filters (CLS8160, Millipore Sigma) and stored at 4°C until use. Antigen degree of labeling was determined by measuring the absorbance at 280, 568, and 646 nm wavelengths to measure total protein, Cy3 dye, and Cy5 dye, respectively. To calculate the concentration of the different constituents, extinction coefficient values of 162000, 271000, 113215, and 141390 M^-1^cm^-1^ were used for one subunit of eOD-60mer, N332-GT2 Trimer, one subunit N332-GT2 Trimer-ferritin (24mer), sulfo-cy3 NHS ester, and sulfo-cy5 NHS ester, respectively. The degree of labeling for either the nanoparticle or soluble antigens was calculated based on the ratio of antigen concentration to Cy3 concentration or to Cy5 concentration.

### Complement binding assay for labeled eOD-60mer

High binding ELISA plates (07-200-37, Fisher Scientific) were coated with 1 μg/mL of labeled and blocked with 2% BSA in PBS. Serum was collected from naïve mice (41.1500.005, Sarstedt), diluted in PBS (3% v/v), and incubated in ELISA plates for 1 hour at 37°C. Complement binding was detected by biotinylated anti-C3 antibody (NB200-540B, Novus Biologicals) followed by streptavidin-HRP (3310-9-1000, Mabtech AB). The plates were developed with 1-Step Ultra TMB-ELISA Substrate Solution (34028, Thermo Fisher Scientific) and the reaction was stopped by adding with 2 N sulfuric acid (BDH7500-1).

### Antigenicity profiling of labeled immunogens

Antigenicity profiles of immunogens following dye conjugation were assessed through binding of structure-sensitive monoclonal antibodies (mAbs) to plate-bound immunogens by ELISA. eOD-60mer with varying amounts of dyes conjugated were directly coated onto Corning High Binding plates (07-200-37, Fisher Scientific) at 2 µg/mL and blocked with 2% bovine serum albumin (A8022, Sigma-Aldrich) dissolved in PBS. For trimers or trimer-NPs, plates were coated with 2 µg/mL *Galanthus Nivalis* Lectin (L8275, Sigma-Aldrich) prior to blocking and capture of 2 µg/mL antigen. Antigenicity profiles were evaluated by adding indicated human monoclonal antibodies at titrated concentrations to plate-bound antigen, followed by detection with mouse anti-human secondary antibody conjugated to HRP (1721050, Bio-Rad Laboratories). HRP binding was determined based on reaction with TMB substrate (34028, Thermo Fisher Scientific) and was stopped by addition of 2 M sulfuric acid (BCH7500-1, VWR) at 1:1 volume ratio. The optical density of the mixture was read out at 450 nm minus the absorbance at 540 nm according to the manufacturer’s instructions.

### *In vitro* antigen digestion assays

Agarose bead-immobilized TPCK-treated trypsin, 150 µL, (20230, Thermo Fisher Scientific) was prepared and equilibrated in 0.1 M ammonium bicarbonate buffer (S8875, Sigma-Aldrich) pH 8 according to the manufacturer’s instructions. These beads were mixed with 100 µL of 20 µg/mL FRET dye-labeled Ags diluted in ammonium bicarbonate buffer and incubated in 37°C. The digested antigens were isolated at specified time points by centrifugation to remove the trypsin beads and characterized using three different assays to identify changes in molecular weight, FRET efficiency, and antigenicity. Molecular weight changes of the antigen were evaluated by reducing SDS-PAGE and protein bands were visualized with high sensitivity Flamingo Fluorescent Protein Gel Stain (1610491, Bio-Rad Laboratories). Intact antigen fractions were determined by digital imaging of gels followed by ImageJ analysis. FRET efficiency and antigenicity changes were obtained by directly coating the recovered antigens onto glass coverslips for imaging or plates for ELISA, respectively.

### Antigen stability within plasma

Blood was collected from naïve mice through retro-orbital bleeding and processed in MiniCollect Tubes with EDTA (450480, MiniCollect) to obtain plasma. To examine antigen stability, 10 μg/mL eOD-60mer_40_ or Trimer_6_ was incubated in diluted plasma (10% v/v in PBS) or 10 μg/mL of porcine trypsin (T4549, Millipore Sigma) at 37°C. The FRET (Ex/Em: 555/665 nm) and Cy5 (Ex/Em: 630/665 nm) signals were recorded at indicated time points using a plate reader and these values were analyzed to determine antigen stability.

### Antigen degradation after incubation with serum

To determine if complement deposition onto antigen prevents enzymatic degradation, 20 μg/mL eOD-60mer was incubated for 15 minutes at 37°C with 10% fresh serum from naïve mice or left alone in PBS. This solution was mixed with trypsin conjugated agarose beads and digested overnight at 37°C. Complement binding and structural integrity of the nanoparticles after digestion was assessed by ELISA using 1 μg/mL anti-C3 and human VRC01 monoclonal antibody, respectively.

### Mouse immunizations

All animal studies were carried out under an IACUC-approved animal protocol following local, state, and NIH guidelines for care and use of animals. C57BL/6J mice were anesthetized and immunized with 10 µg of indicated antigens in the presence or absence of 5 µg saponin adjuvant subcutaneously at the left and right sides of the tail base.

### Tissue processing

Inguinal lymph nodes extracted from euthanized mice were submerged into cryomolds containing O.C.T. (23-730-571, Fisher Scientific) compound and dipped into 2-methylbutane (M32631, Millipore Sigma) pre-chilled in liquid nitrogen for 5-10 minutes prior. All frozen tissues were cryosectioned on a Leica CM1950 at 10 µm thickness, adhered to Superfrost Plus microscope slides (12-550-15, Fisher Scientific), and stored in -80°C until use.

### FRET imaging and analysis

To assess antigen integrity, an acceptor photobleaching method was used to compared Cy3 dye emission intensity before and after photobleaching the Cy5 dye. FRET imaging was carried out on a Leica SP8 Confocal Microscope with a 25x objective, imaging selected square regions with dimensions ranging from 175 to 250 μm. Cy3 signal was recorded by exciting the dye at 555 nm and collecting its emission between 565 and 615 nm wavelengths, while Cy5 signal was recorded after exciting the dye at 640 nm and collecting emission between 660 and 720 nm wavelengths. To photobleach the Cy5 dye, regions of interest in lymph node sections were excited with 640 nm wavelengths lasers at maximum power until the Cy5 intensity was less than 10% of its pre-bleached values; the duration of the bleaching process varied between 90 and 180 seconds. Cy3 and Cy5 fluorescence signals were then collected again post-acceptor bleaching. FRET efficiencies (*E*) of antigens were determined based on established methods (Roszik et al., 2008). FRET efficiency *E* at each pixel (*E*_*ij*_) is given by:

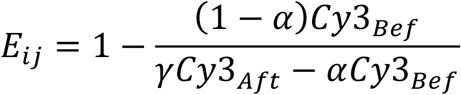

where *Cy*3_*Bef*_ and *Cy*3_*Aft*_ are Cy3 emissions before and after Cy5 photobleaching. Correction factors *α* and *γ* account for incomplete bleaching of the Cy5 (0 ≤ *α* ≤ 1) and unintended photobleaching of Cy3 (*γ*≥ 1), respectively. *α* values were experimentally determined from the ratio of Cy5 emission signals before and after photobleaching, while *γ* was determined to be ∼1.05 based on coverslip-coated antigens conjugated only with Cy3. Moreover, correction factors, *δ*, crosstalk from acceptor emission into donor emission channel, and *ε*, crosstalk of photodegraded acceptor product into donor emission channel, that are listed in the full equation for *E* are found to be 0.004 ± 0.00094 and 0.0062 ± 0.00077, respectively, from monolayer experiments with antigens labeled only with Cy5. Thus, both *δ* and *ε* were approximated to be zero. To quantify *E*, antigen locations were first identified by using a binary mask generated from thresholding intensity values in the Cy5 image prior to photobleaching. To this end, non-immunized LNs were imaged under same conditions to approximate the background fluorescence value of Cy5 needed for thresholding. This mask was applied to the remaining images to quantify *Cy*3_*Bef*_ from the Cy3 image before photobleaching, *Cy*3_*Aft*_ from the Cy3 image after photobleaching, and *α* from the Cy5 images before and after photobleaching. These parameters were used to generate a histogram of *E* values for antigens found within each imaging area. To determine the fraction of degraded antigen, this procedure was repeated for an intact control sample comprised of FRET dye-conjugated antigen coated onto a glass coverslip. To quantify the amount of fully intact antigen at each time point/condition was determined by quantify the fraction of antigen^+^ pixels in each imaged region with *E*_*ij*_ values larger than the minimal *E* value detected in the intact control sample.

For examining fractions of intact antigens co-localized with specified protein markers, the acceptor photobleaching procedure was carried out on LN sections that were fixed, immunostained with antibodies, and mounted in PBS. In addition to the 4 images required for calculating *E*, fluorescence images of cell markers were recorded and overlaid onto regions of intact and degraded antigens.

### Vibratome sectioning

Low melting point agarose (A4018, Sigma Aldrich) solution was prepared by adding 0.4 g of agarose into 20 mL of PBS to achieve a concentration of 2% wt/v. This mixture was microwaved at 30 seconds intervals until the agarose was fully dissolved and the solution was equilibrated at 37 °C for at least 45 minutes. Inguinal LNs were harvested from immunized or naïve mice and placed onto a 35 mm diameter Petri dish. The agarose solution was carefully added to this dish and incubated at 4 °C for 10 minutes. A rectangular block encompassing the LNs were excised and mounted onto the sample holder of a Leica Vibratome VT1200s using Vetbond tissue adhesive (3M). The sample was held in PBS pre-chilled to 4°C and sectioned to obtain 250 μm thick live tissue sections that were immediately transferred to a petri dish containing ice cold RPMI 1640. All slices were collected and maintained in this solution until further use.

### In vitro AZP Assay

The sequences for the custom peptide (CPC Scientific) probes are shown below (Soleimany et al., 2021a).

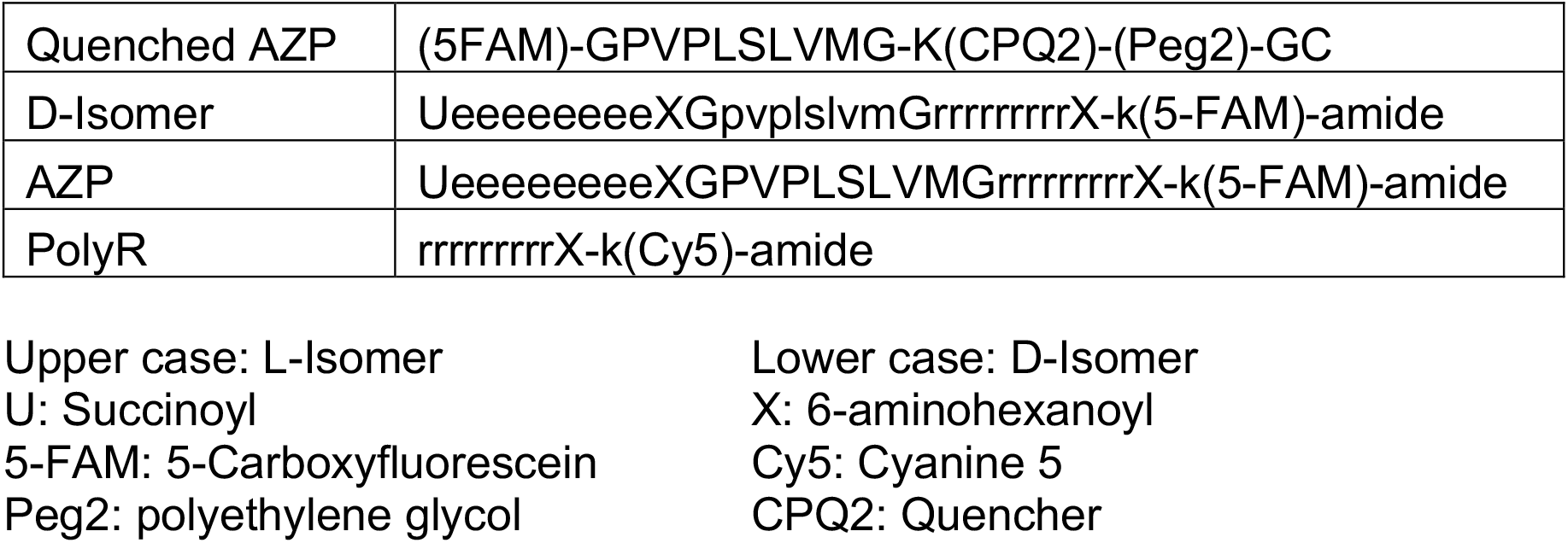

To analyze the cleavage of peptide sequence in the AZP probe by different recombinant proteases, a quenched version of the probe that consisted of a peptide sequence flanked by quencher CPQ2 and fluorophore FAM was used. This reaction was carried out by incubating 1 μM of the quenched AZP probe with 1 μg/mL of recombinant MMP14 (918-MP-010, R&D Systems), ADAM17 (2978-AD-10, R&D Systems), ADAM10 (946-AD-020, R&D Systems), Cathepsin D (1029-AS-010, R&D Systems), Cathepsin S (50769-M08H, Sinobiologicals), or Cathepsin L (1515-CY-010, R&D Systems). The buffer pH for digestion was adjusted based on manufacturer’s protocol and were 8.5, 9.0, 9.0, 3.5, 4.5, and 6.0 for MMP14, ADAM17, ADAM10, Cathepsin D, Cathepsin S, and Cathepsin L, respectively. The FAM signal (Ex/Em: 485/535 nm) was recorded on a plate reader initially and after 60 minutes of incubation at 37°C.

For visualizing protease activity within tissues, LNs were harvested from naïve mice or mice immunized with eOD-60mer and saponin at indicated time points. These tissues were embedded within 2% agarose gel containing 5 µM AZP and PolyR control probes, vibratome sliced to obtain 250 µm thick live sections, and maintained within RMPI 1640 supplemented with 2 µM ZnCl_2_. Parallel tissue sections were also incubated in 5 µM uncleavable D-Isomer probe and PolyR as a control. After 2 hours incubation at 37°C, the tissue-gel constructs were washed twice with PBS before fixation in 4% paraformaldehyde (PFA) overnight at 4°C. Excess PFA was removed by washing in PBS and these sections were imaged directly by confocal microscopy. From the images, areas of positive AZP probe signal normalized to areas with positive PolyR probe signal was calculated for inside versus outside of the follicles.

To immunophenotype proteolytically active cells, live LN vibratome sections were placed into RMPI medium containing the following combination of probes at 2 µM concentration: (i) AZP and PolyR probes, (ii) D-Isomer and PolyR probe, and (iii) AZP probe only. These slices were maintained in the probe media for 2 hours at 37°C prior to washing away the excess probe by performing two cycles of transferring the slices to and maintaining in fresh PBS for 15 minutes at 37°C. The LN sections were subsequently processed for flow cytometry analysis. To assess protease activity, different cellular phenotypes were first identified and AZP^+^ cells were determined only amongst the PolyR^+^ cells within each cell type. Gating for PolyR and AZP probe binding was determined based on cells extracted from slices incubated in probe media conditions (iii) and (ii), respectively (**Figure S5A**).

### Ex vivo modulation of protease activity within LN slices

To examine the impact of proteases on FDC-localized antigens, mice were first immunized with saponin and eOD-60mer_40_. After 3 days, LNs were harvested and vibratome sliced to 250 µm thickness and were left untreated in RPMI1640 medium or exposed to either 200 µg/mL porcine trypsin (T4549, Millipore Sigma) or 10 µg/mL recombinant full length murine MMP9 without additional chemical activation by p-Aminophenylmercuric acetate (ab39309, Abcam). After 4 hours at 37°C, tissue slices were washed in PBS and fixed overnight in 10% Neutral Buffered Formalin (HT5011-1CS, Sigma-Aldrich) at 4°C prior to confocal imaging.

Similarly, the effect of protease inhibitors on antigen degradation was assessed by immunizing mice with saponin and eOD-60mer_40_, followed by harvesting of LNs 2 hours later for vibratome sectioning. These live slices were left untreated in RPMI1640 medium or incubated with 100 µg/mL of Marimastat (M2699, Millipore Sigma) or Halt Broad-spectrum protease inhibitor without EDTA (78430, Thermo Fisher Scientific) at a five-fold dilution of the stock solution. After 6 hours in 37°C, the tissue slices were washed in PBS and fixed overnight in 10% Neutral Buffered Formalin (NBF) at 4°C.

In both *ex vivo* experiments, the fixed sections were incubated in fresh PBS for 1 hour at 25°C followed by flash freezing and cryosectioning. The sections adhered to microscope slides were mounted in PBS and imaged immediately.

### Single-cell RNA-sequencing analysis

The scRNA-seq data was collected and analyzed in a separate study (Cui et al. in progress). The data was used here to identify cell type-specific expression of proteases. Briefly, scRNA-seq (10X Genomics) was performed according to manufacturer’s instructions on dissociated cells from lymph nodes of mice that were either untreated or 6 hours following vaccination with 200 µg OVA peptide and 20 µg CpG 1826 adjuvant. Cell Ranger was used to process sequence data. Gene expression is log-normalized for each cell such that the total unit for each cell sums to 10,000. Clusters of the major cell types were derived using the Louvain clustering algorithm and annotated based on known markers. Genes encoding extracellular or secreted proteases, including ADAMs, ADAMTSs, MMPs and elastases, were included in the analysis.

### Immunofluorescence staining

Frozen sections were retrieved from -80°C, quickly thawed, and incubated in 10% neutral buffered formalin solution for 8.5 minutes at 25°C. The sections were washed 3 times in PBS with 10-minute incubation time between each wash. Excess PBS was removed after the final wash before incubating the slides in blocking buffer, comprised of 2% bovine serum albumin (BSA) and 2% Triton X-100 in PBS. After 30 minutes, the blocking buffer was aspirated and the slides were stained in the following primary antibody solutions also made in blocking buffer for ∼16 hrs at 4°C: 1:75 anti-ADAM17 (NBP2-67179, Novus Biologicals), 1:75 anti-ADAM10 (NBP176973, Novus Biologicals), 1:75 anti-MMP14 (ab51074, Abcam), 1:75 anti-CD35 (740029, BD Biosciences), 1:75 anti-LYVE1 (53-0443-82, Thermo Fisher Scientific), 1:75 anti-CD169 (142421, BioLegend). These slides were washed in PBS 3 times for 10 min, stained with 1:200 diluted secondary antibodies solution in blocking buffer (ab150063 Abcam, ab150061, Abcam) for 4 hours at room temperature, and washed again in PBS. To mount the slides, 5 µL of PBS was added directly onto the stained tissues prior to gently placing a 20×20 mm coverslip on top of the PBS droplet to sandwich the section. The coverslip was sealed using CoverGrip coverslip sealant (23005, Biotium) and imaged immediately. Slides were stored at 4°C for maximum of 1 week.

For VRC01 mAb staining, slides that were fixed, washed in PBS, and incubated either in blocking buffer or in 2% BSA-PBS solution. For permeabilized sections in blocking buffer, slides were stained with human VRC01 mAb directly labeled with AF488 NHS ester (21820, Lumiprobe). For non-permeabilized sections, slides were first stained with biotinylated anti-Cy3/Cy5 antibody (C3117, Millipore Sigma) at 1:100 dilution in 2% BSA-PBS prior to using streptavidin BV421 (405225, BioLegend) and anti-human VRC01 mAb conjugated to AF488.

### Human tonsil tissue acquisition and processing

Adult human tonsil samples sourced from standard surgical procedures performed for clinical care was provided by the Massachusetts Eye and Ear Infirmary at the Massachusetts General Hospital. Usage for scientific research was approved by the Massachusetts General Hospital Institutional Review Board under Protocol P2020-003521. Tonsil samples were deidentified and processed within 2–3 h after removal. Acquired tonsil tissues were flash frozen, cryosectioned, and immunostained in the same manner as murine lymph nodes.

### Confocal Microscopy

For all experiments, imaging was performed on a Leica SP8 confocal microscope equipped with a white light laser and spectral emission filter to detect emission wavelengths between 470 and 670 nm with a minimum bandwidth of 10 nm. All images were recorded with a 25X water immersion lens and for assessing antigen drainage in the LNs, laser settings were kept constant across different time points for each immunogen.

### Flow Cytometry

For analysis of AZP binding and protease expression, inguinal lymph nodes from immunized or naïve mice or live LN sections were incubated in 1 mL of digestion solution comprised of RPMI1640 medium containing 0.8 U/mL Dispase (07913, Stemcell Technologies), 0.1 mg/mL Collagenase D (11088858001, Sigma-Aldrich), and 0.1 mg/mL of DNAse (10104159001, Sigma-Aldrich). After 20 minutes incubation at 37 °C, the LNs were repeatedly passed through a 1 mL pipette tip to mechanically rupture the tissue. Care was taken to collect ∼ 1 mL of the solution containing only the liberated cells prior to transferring to ice cold PBS containing 5 mM EDTA and 2% fetal bovine serum (stop buffer). The remaining tissue debris was further processed by a second incubation with 1 mL of fresh digest buffer for 20 minutes at 37 °C and transfer of the supernatant to stop buffer. This solution was passed through a 40 µm pore size filter, centrifuged, and transferred into a V-bottom 96 well plate for flow staining. For germinal center analysis, cells were isolated by mechanically crushing the LNs against a filter cap with 70 µm pore size and generously pipetting cold PBS containing 1% BSA and 5 mM EDTA (FACS buffer) onto the porous mesh. The cells were centrifuged, resuspended in flow cytometry buffer, passed through a 40 µm pore size filter, and transferred to a V-bottom 96 well plate.

The isolated cells were stained with Live/Dead fixable aqua stain (L34957, Thermo Fisher Scientific) for 10 minutes at 25°C prior to washing twice in flow cytometry buffer. Cells were then incubated with Fc Block for 10 minutes at 4°C before staining with antibodies listed in the Key Resource table for 20 additional minutes at 4°C. For on-target trimer and off-target probe specific GC B cell analysis, cells stained with antibodies were distributed evenly and exposed to biotinylated trimer or peptide probes preincubated with streptavidin (30 minutes at molar ratio of 1:4 at 25°C) conjugated to PE (405203, BioLegend), BV421 (405226, BioLegend), or APC (40520L7, BioLegend). For AZP analysis, the stained cells were immediately analyzed (Soleimany et al., 2021b). For protease expression analysis, cells stained with surface markers were fixed and permeabilized using transcription factor staining buffer kit (00-5523-00, Invitrogen) followed by incubation with anti-protease antibody for 30 minutes at 25°C. After washing twice with wash buffer, the cells were incubated with donkey anti-rabbit antibody conjugated to Alexa Fluor 488 dye to detect the proteases. For compensation, UltraComp eBeads^™^ Plus compensation beads (01-3333-42, Thermo Fisher Scientific) were used to bind to dye-labeled antibodies. Stained cells were analyzed using a BD FACSymphony A3.

### Probe synthesis for characterizing Trimer Breakdown product specific GC B cells

Peptides corresponding to and buried within BG505 MD39 were synthesized by GenScript, Inc. Sequences of the peptides are as follows:

HR1 peptide-biotin, GIKQLQARVLAVEHYLGAGK-biotin;

C1 peptide-biotin, DQSLKPAVKLTPLGAGK-biotin;

V3 peptide-biotin, TRPNNNTVKSIRIGPGQAFYYTGGAGK-biotin.

### ELISA analysis of serum antibody responses

Blood collected from immunized mice through retro-orbital bleeding was processed in collection tube with separator gel (41.1500.005, Sarstedt) to obtain serum samples. To analyze on-target antibody response, high binding ELISA plates (07-200-37, Fisher Scientific) were coated with 1 μg/mL trimer and blocked with 2% BSA in PBS. For off-target responses, plates were first coated with 2 μg/mL streptavidin (434302, Thermo Fisher Scientific), blocked with 2% BSA in PBS, and incubated with 2 μg/mL of HR1, C1, or V3 peptides diluted in blocking buffer. To detect antigen-specific IgG responses, dilutions of serum were added to the wells and incubated for 1.5 hours at 25°C. Plates were washed 3 times in PBS containing 0.2% Tween-20, then anti-mouse IgG secondary antibody conjugated to HRP (172-1011, Bio-Rad Laboratories), diluted 1:5000 in blocking buffer as per manufacturer instructions, was added to the wells. After 1 hour incubation, plates were again washed, developed with TMB, and stopped with sulfuric acid. Endpoint titers are reported as inverse dilutions where absorbance at 450 nm minus the reference absorbance at 540 nm equals 0.1.

## QUANTIFICATION AND STATISTICAL ANALYSIS

Information pertaining to statistical analysis such as n (number of mice or tissue slices or image regions) and p values are indicated in the figure legends. All plots within the figures show mean. No methods were used to determine if data met assumptions of statistical approach. Graphpad Prism 9 was used to calculate statistical significance from tests specified within each figure legend. Statistically significant differences were considered to be at *p ≤ 0.05, **p ≤ 0.01, ***p ≤ 0.001, and ****p ≤ 0.0001.

