## Supplementary Information for "Spatially regulated protease activity in lymph nodes renders B cell follicles a sanctuary for retention of intact antigens"

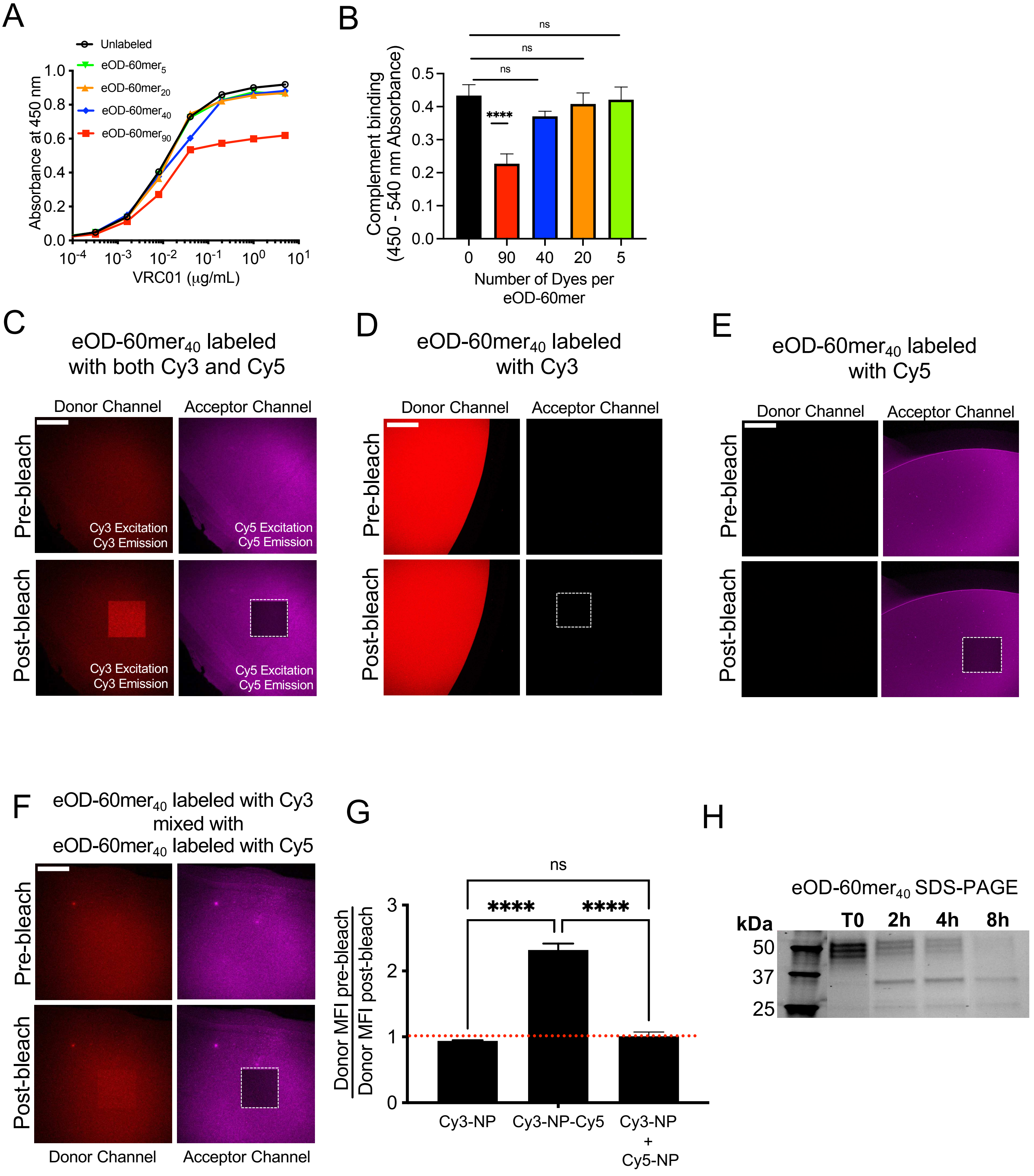


Supplementary Figure 1

**Figure S1: Characterization of dye-labeled eOD-60mer and FRET analysis, Related to Figure 1.**

(A) ELISA absorbance of VRC01 mAb binding to eOD-60mer conjugated with specified number of dyes (subscript) coated on plates. eOD-$60mer_{40}$ indicates approximately 20 dyes of Cy3 and Cy5 dyes on each nanoparticle.

(B) ELISA analysis of C3 complement binding to plate-bound eOD-60mer conjugated with different amounts of dyes following incubation of 60mer particles with 10% mouse serum for 1 hr at 37°C (n=3 samples/group). ns, not significant; ****, $p \leq 0.0001$ by one-way ANOVA with post-hoc Dunnett test for pair-wise comparison against unlabeled control 60mer.

(C-F) eOD-60mer conjugated with specified dyes was coated onto a glass coverslip and imaged by confocal microscopy. Donor channel (Cy3 Ex/Em) and Acceptor Channel (Cy5 Ex/Em) are shown before and after photobleaching Cy5. Dashed region in lower right panel indicates the region that is photobleached.

(G) Quantification of the change in Cy3 (Donor) emission after photobleaching for eOD-60mer (NP) conjugated with different dyes. (n=5-6 regions imaged from 2 independent experiments. ns, not significant; ****, $p \leq 0.0001$ by one-way ANOVA with post-hoc Tukey test for multiple pair-wise comparisons,

(H) SDS-PAGE gel image of eOD-$60mer_{40}$ digested with trypsin beads for the indicated times.

All scale bars indicate 100 μm in length. All graphs show mean±s.d.

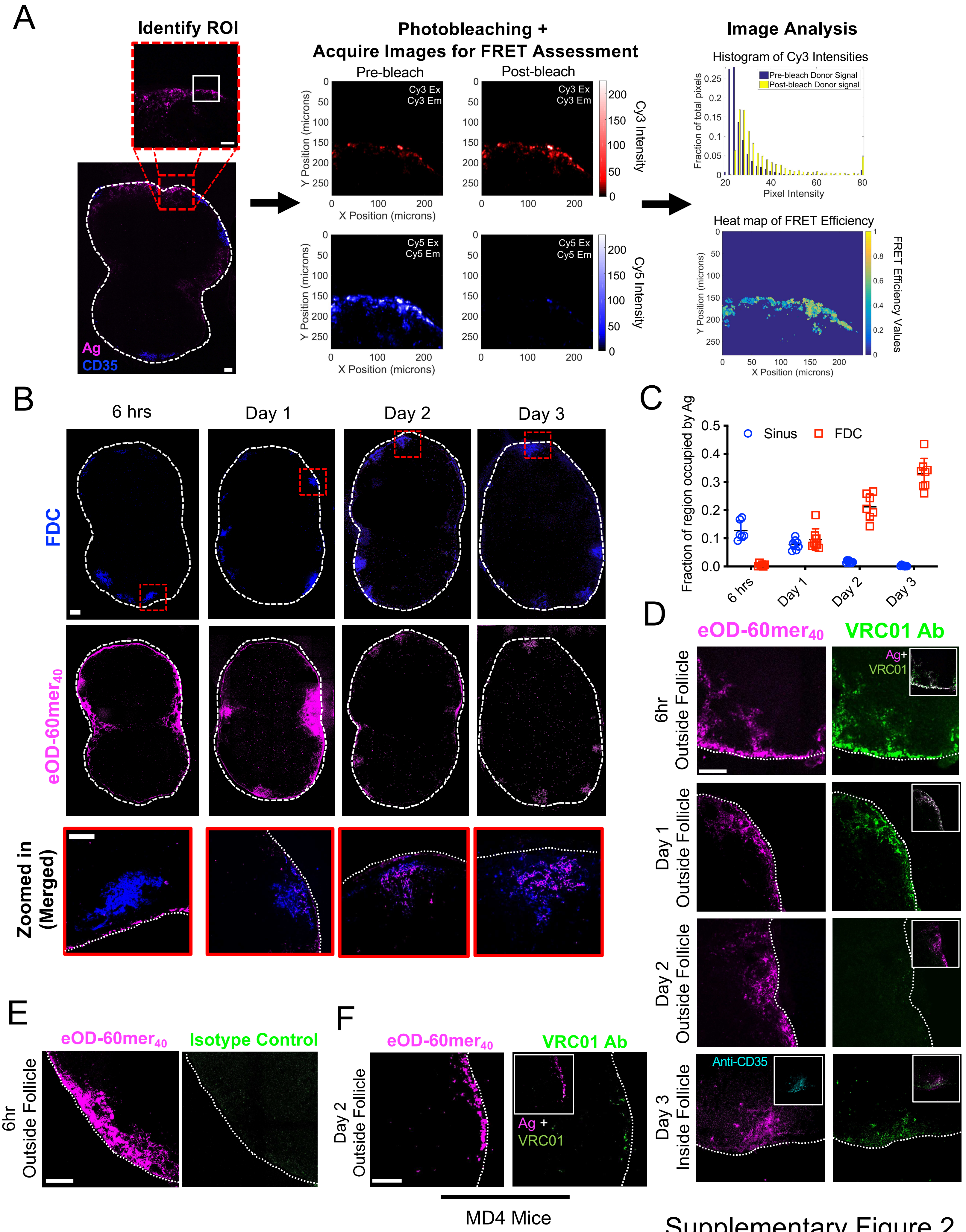


Supplementary Figure 2

**Figure S2: Antigen trafficking and stability analysis, Related to Figure 1 and 2.**

(A) Workflow for FRET imaging of tissue sections containing eOD-60mer$_{40}$ (magenta) and stained with anti-CD35 antibody (blue) to label the FDC network. Antigen is first identified using Cy5 Excitation/Emission and a set of donor and acceptor images are recorded before and after photobleaching. Histogram shows the change in Cy3 intensity distribution which is used to calculate FRET Efficiency (*see* Materials and Methods, FRET Imaging and Analysis).

(B-E) Groups of C57BL/6J mice (n=3/group) were immunized with 5 μg saponin adjuvant and 10 μg eOD-60mer$_{40}$. At indicated time points, LNs were harvested, flash frozen, and sectioned for confocal imaging.

(B) Representative confocal images of LN sections containing eOD-60mer$_{40}$ (magenta) and stained with anti-CD35 antibody (blue) at different time points. Bottom panel shows magnified image of region within the window with red dashed line.

(C) Quantification of antigen localization within subcapsular sinus (Sinus) and FDC. Each point represents a tissue section. Data collected from at least 6 tissue sections from 6 LNs. Shown are mean±s.d.

(D) Representative regions within fixed and permeabilized sections of LNs harvested at indicated time after immunization. These sections contain eOD-60mer$_{40}$ (left) and were stained with labeled VRC01 Ab (right). Insets show merged images of the antigen and labeled VRC01 mAb.

(E) Representative region within fixed and permeabilized sections of LNs harvested 6 hrs after immunization. The sections contain eOD-60mer$_{40}$ (left) and stained with labeled isotype control for VRC01 Ab (right).

(F) Groups of MD4 mice (n=2/group) were immunized as in (B). After two days, 4 inguinal LNs were harvested, flash frozen, and sectioned for confocal imaging. Representative regions within fixed and permeabilized sections of LNs containing eOD-60mer$_{40}$ (left) and stained with labeled VRC01 Ab (right). Insets show merged images of the antigen and labeled VRC01 mAb.

White dashed lines in all of the confocal images indicate the contour of the LN sections.

All scale bars indicate 100 μm in length.

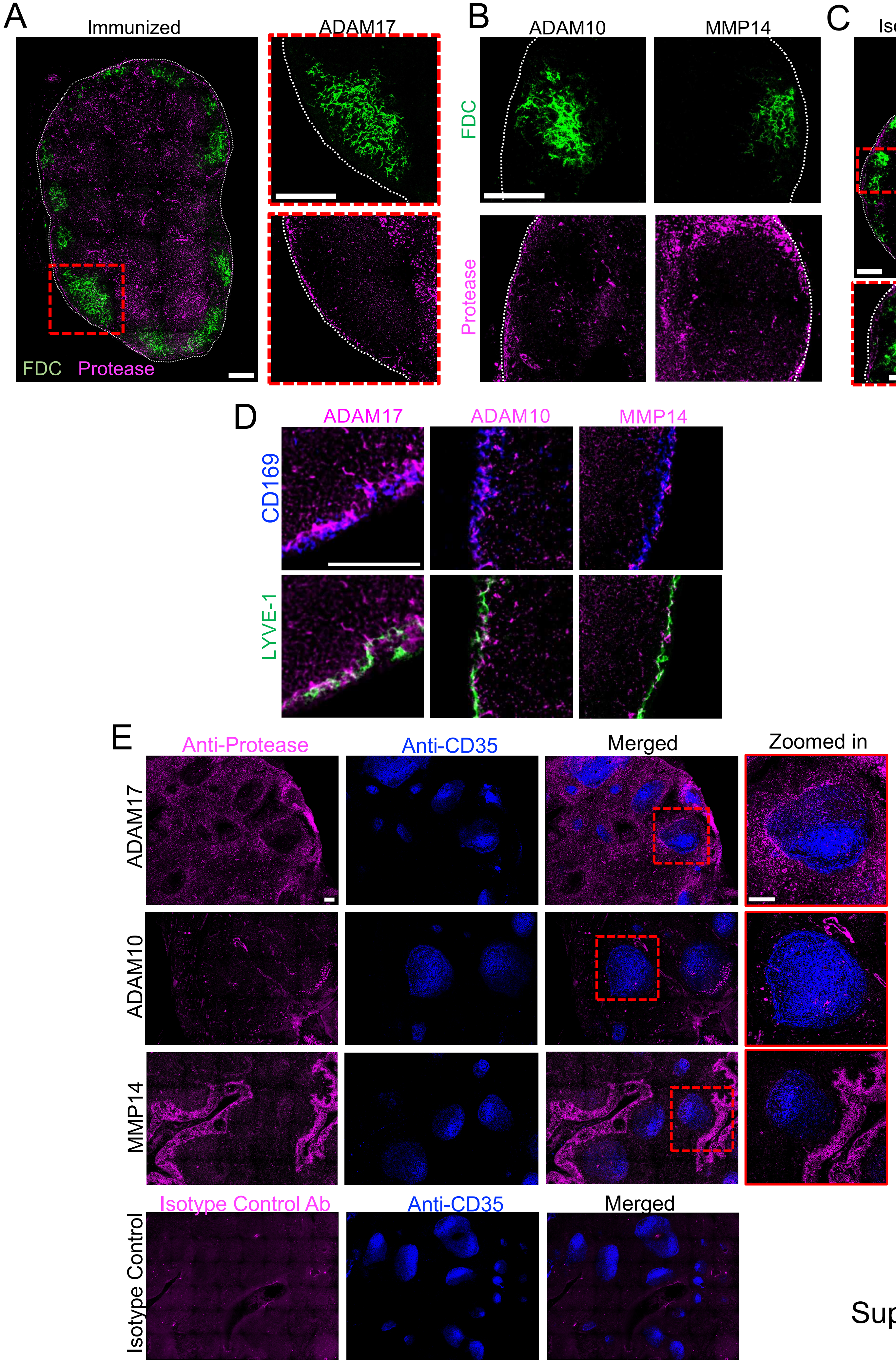


Supplementary Figure 3

**Figure S3: Metalloproteinase expression within lymphoid tissues, Related to Figure 3.**
(A-B) LNs from C57BL/6J mice immunized with 5 µg saponin adjuvant and 10 µg eOD-60mer were harvested after 1 day for tissue sectioning and staining. White dash lines indicate the periphery of the LN section in the following confocal images.
(A) Tissue section stained with anti-ADAM17 and anti-CD35 antibodies. The magnified images on the right shows the FDC network or proteases within the red dashed region identified in the lower magnification view. Scale bars: 200 $\mu$m.
(B) Magnified images around the follicle region showing the FDC network and indicated proteases. White contour encircles the B-cell follicle. Scale bars: 200 $\mu$m.
(C) LN sections from naïve C57BL/6J mice stained with an isotype control (magenta) and anti-CD35 antibody staining (green). Magnified image within the red dashed region is shown on the bottom. Scale bars: 200 $\mu$m.
(D) Lymph nodes were harvested from naïve C57BL/6J mice (n=3/group), sectioned and stained with antibodies against indicated proteins. Representative regions near the subcapsular sinus showing the colocalization of indicated proteases with $CD169^+$ macrophages (Macs) or LYVE-$1^+$ LECs. Scale bars: 100 $\mu$m.
(E) Representative sections of human tonsil sections stained with antibodies against the indicated proteases (magenta) or isotype control antibody (magenta, bottom row) and anti-CD35 staining (blue). Magnified images of regions within red dashed regions are shown on the right. Scale bars: 200 $\mu$m.

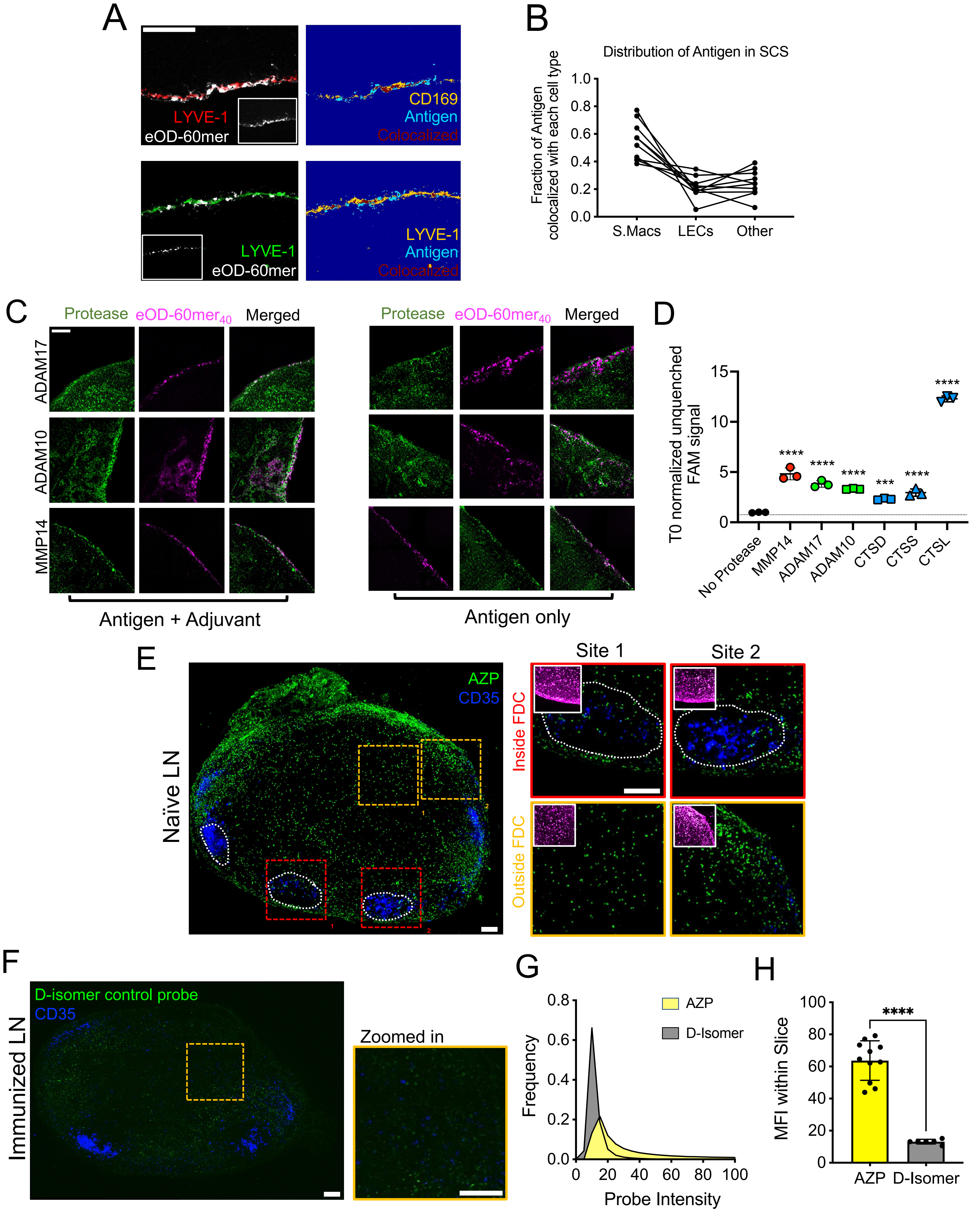


Supplementary Figure 4

**Figure S4: Visualizing proteolytically active cells within lymphoid tissues, Related to Figure 3 and 4.**

(A-C) C57BL/6J mice (*n*=3 animals/group) were injected with 10 µg eOD-60mer$_{40}$, and 6 hours later draining LNs were harvested, sectioned and stained with antibodies against the indicated proteins.

(A) Representative image of stained subcapsular macrophages (CD169$^{+}$) and LECs (LYVE-1$^{+}$) in the SCS (left). Inset shows eOD60mer$_{40}$ signal. Processed images show co-localization of antigen with each cell type (right).

(B) Quantification of protease colocalization with different cell types. Each point represents a region along the sinus and connected line indicates cells analyzed from same region of interest. Data collected from 6 sections from 6 different lymph nodes.

(C) Lymph node sections from C57BL/6J mice (n=3/group) injected with 10 µg of eOD-60mer$_{40}$ for 6 hours alone or together with 5 µg of saponin adjuvant 24 hours prior. These sections were also stained for indicated proteases (green).

(D) *In vitro* cleavage of 1 µM quenched AZP probe in the absence of proteases (No Protease) or after incubation with 1 µg/mL recombinant MMP14, ADAM17, ADAM10, Cathepsin D (CTSD), Cathepsin S (CTSS), or Cathepsin L (CTSL) for 1 hr at 37°C. FAM signal increases upon cleavage of the probe. Shown are fluorescence values normalized to a value of 1 for the uncleaved probe. (n = 3 samples/group). ****, $p \leq 0.0001$; ***, $p \leq 0.001$ by one-way ANOVA with post-hoc Dunnett test for pair-wise comparison against No Protease.

(E) Lymph nodes from naive C57BL/6J mice (n=3 animals/group) were harvested and prepared as in Fig. 4B. Representative whole-LN tissue section (left) and higher-magnification images (right) taken from dashed regions “inside FDCs” (red dashed regions) or “outside FDCs” (orange dashed regions). Main images show AZP probe (green) overlaid on anti-CD35 staining (blue), while insets at right show the corresponding staining from the same areas for the control probes (magenta). Regions within dashed white lines indicate follicles.

(F-H) Naive C57BL/6J mice (n=3/group) were prepared as in (E) except the slices were incubated in uncleavable D-Isomer control probe.

(F) Representative whole-LN tissue section and magnified image of dashed region are shown to the right.

(G-H) Representative probe intensity distribution (G) and MFI (H) for AZP or D-Isomer treated slices. Each point in (H) represents a single slice. Data collected from at least 6 slices across 6 LNs. (T-test comparison between AZP and D-isomer MFI, ****$p \leq 0.0001$).

All scale bars indicate 100 µm in length. All graphs show mean±s.d.

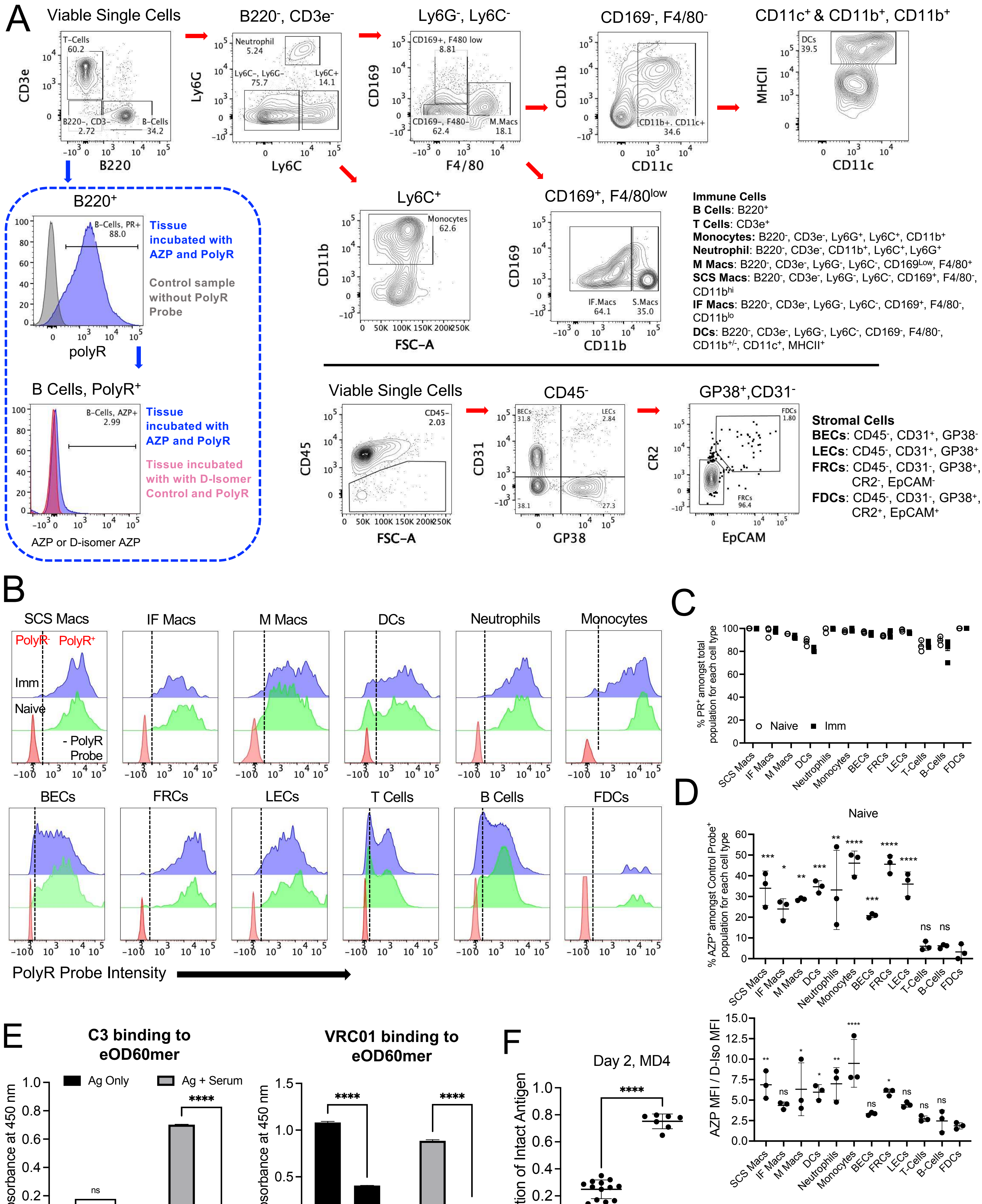

Supplementary Figure 5

**Figure S5: Identification of proteolytically active cell populations, Related to Figures 3 and 4.**

(A-D) C57BL/6J mice (n=3 animals/group) were left naïve or immunized with 10 µg of eOD-60mer and 5 µg saponin adjuvant. 24 h later, LNs were harvested and vibratome sectioned into 250 µm thick live tissue slices. Tissue slices were incubated in AZP or D-Isomer Probe together with PolyR control probe for 2 hours at 37°C, followed by flow cytometry analysis of probe binding.

(A) Gating strategy to identify LN cell populations and proteolytically active cells using flow cytometry. Top and bottom panels separated by horizontal line show gating strategies for lymphocytes and stromal cells, respectively. Red arrows indicate the hierarchical gating path. Blue arrows illustrate how proteolytically active cell fraction was determined for each cell type using B cells as an example. Specific markers used to identify different populations are also listed.

(B) Representative histogram showing PolyR binding amongst cells from naïve LN slices (Naive), immunized LN slices (Imm), and naive LN slices that are not treated with PolyR probe (- PolyR Probe). Dashed line denotes threshold defining PolyR positive ($PR^+$) and negative ($PR^-$) cells.

(C) Quantification of fraction of cells bound to PolyR probe amongst different cell types in naïve and immunized LNs.

(D) Quantification of fraction of cells bound to AZP probe (top graph) and ratio of AZP probe to D-Isomer probe MFI (bottom graph) amongst different cell types in naïve LNs. (One-way ANOVA with post-hoc Dunnett test for pair-wise comparison against FDC, ****$p \le 0.0001$, ***$p \le 0.001$, **$p \le 0.01$, *$p \le 0.05$).

(E) eOD-60mer pre-incubated with 10% fresh serum from naïve mice and was left intact or digested with trypsin conjugated agarose beads for 16 hrs at 37°C. The treated nanoparticles were then coated on plates and ELISA analysis was carried out to detect complement C3 deposition on the particles (left graph) and to assess the degree of antigen integrity via VRC01 binding (right graph). (n = 3 samples /group, T-test comparison between intact and digested samples within each group, ****$p \le 0.0001$).

(F) Groups of MD4 mice (n=2 animals/group) were immunized with 10 µg of eOD-$60mer_{40}$ and 5 µg saponin adjuvant. After two days, inguinal LNs were harvested, flash frozen, and sectioned for imaging. FRET analysis of dye-labeled antigen outside and inside the FDC. Data collected from at least 6 tissue sections from 4 LNs. (T-test comparison between outside and inside follicles, ****$p \le 0.0001$).

All graphs show mean±s.d.

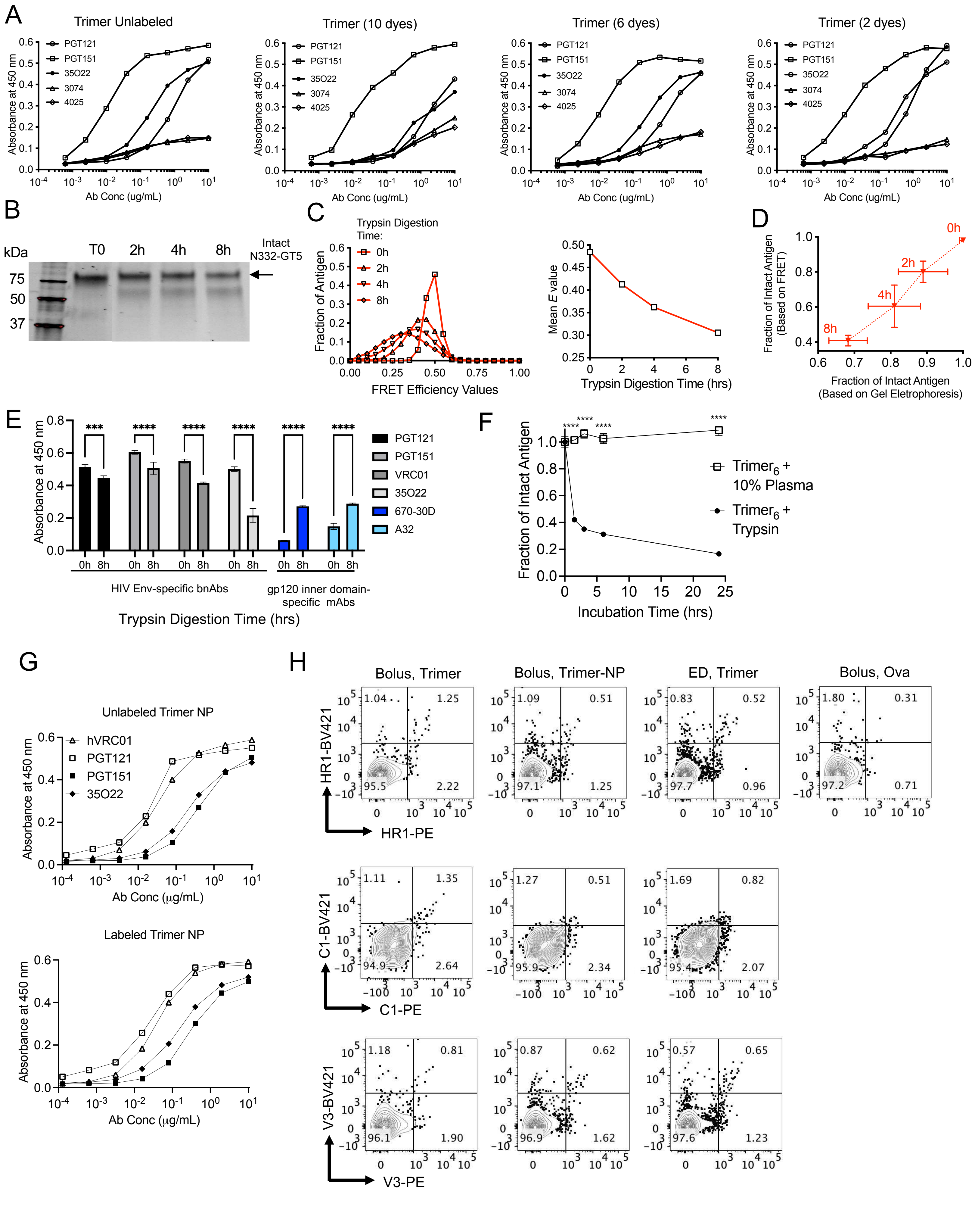


Supplementary Figure 6

**Figure S6: Characterization of dye-labeled HIV Env trimers and trimer-induced humoral immune responses, Related to Figures 5 and 6.**

(A) ELISA analysis of labeled trimer antigenicity profiles, as characterized by structure-sensitive mAb binding after conjugation with indicated dye quantities. Trimer (10 dyes) indicates ~5 Cy3 and 5 Cy5 dyes conjugated onto the protein structure.

(B) SDS-Page characterization of $MD39_6$ molecular weight change after digestion with agarose bead-immobilized trypsin for specified times.

(C) Histograms of FRET Efficiency values collected from $MD39_6$ treated with trypsin for indicated times and then coated on glass coverslips for imaging. Inset shows mean E values at each time point.

(D) Comparison of measured proportion of intact eOD-$60mer_{40}$ as measured by FRET vs. SDS-PAGE (n=3samples/time point analyzed by FRET and SDS-PAGE).

(E) Comparison of structure-sensitive mAb (blue) binding to $MD39_6$ following trypsin digestion for 8 hours. Binding of mAbs that recognize epitopes on the interior of unfolded trimers (shades of blue) were also used to assess trypsin mediated degradation. (n = 3 samples/time point). ****, $p \leq 0.0001$ by t-test comparison between 0 and 8 hours incubation time for each antibody.

(F) Plate reader analysis of 10 μg/mL $Trimer_6$ incubated with plasma from naïve C57BL/6J mice diluted to 10% v/v in PBS or 10 μg/mL of Trypsin at 37°C for specified amount of time. Time 0 normalized FRET signal (Cy3 excitation and Cy5 emission) divided by Cy5 signal (Cy5 excitation and Cy5 emission) is shown at each time point. Graph shows mean±s.d. (n = 3/group, T-test comparison between 60mer mixed with diluted plasma or Trypsin, ****$p \leq 0.0001$).

(G) ELISA analysis of changes in Trimer-Ferritin (Trimer NP) antigenicity profiles, as assessed by structure-sensitive mAb binding after conjugation with 20 Cy3 and 20 Cy5 dyes.

(H) Representative flow cytometry plots showing proportions of GC B cells specific for off-target probes shown in Figure 6C. This analysis was carried out 14 days after injection of the initial dose.
